# Rapid Emergence of an Adaptive Spatial Representation

**DOI:** 10.1101/2024.09.13.612867

**Authors:** Meike M. Rogalla, Hannah L. Nemeth, Clara Martinez-Voigt, Gunnar L. Quass, Alexander N. Ford, Gunseli Wallace, Gabriel Corfas, Pierre F. Apostolides

## Abstract

Peripheral sensory cue availability fluctuates across timescales ranging from seconds to years. This requires central circuits to continuously adapt their representations while preserving established codes, but how the brain achieves this flexibility is unknown. Using two-photon Ca^2+^ imaging in the mouse inferior colliculus, we address this knowledge gap by testing how neurons encode sound location before and after abruptly disrupting binaural cues via monaural hearing loss. Despite measurable single neuron changes, altering binaural cues had remarkably little impact on the population’s ability to encode auditory space. Rather, neural activity rotated into a spatially informative representational subspace, orthogonal to the original binaural axis, within two hours of hearing loss. This rotation occurred in absence of overt experience or training, and reflected a shift from ON- to OFF-dominated spatial selectivity. Thus, central population codes rapidly reorganize to accommodate sudden changes at the periphery, without overwriting existing representations.

## 1 Introduction

Navigating the environment depends on internal neural representations of external space, which emerge from the coordinated activity of neuronal populations ^1–4^. Hearing is particularly important in spatial processing: the location and movement of distant or visually obscured objects can be inferred from sound alone ^5–8^, conferring a survival advantage to predator and prey alike. Unlike vision or touch, however, the auditory periphery contains no explicit spatial map. Sound location is instead computed centrally by integrating binaural cues such as interaural timing and level differences, and spectral cues such as the direction-dependent filtering of sound frequencies by the pinnae ^9–12^. For such computations to be useful, sound localization circuits must remain flexible in the face of continuous changes in cue reliability. For example, ambient reverberation shifts abruptly when one walks from a carpeted hallway into a tiled lecture hall, a child’s head and body grow continuously across development, and an adult’s peripheral sensitivity declines during aging. How the auditory system flexibly adapts to shifting cue availability across multiple time scales is unknown.

Monaural hearing loss provides an extreme test of this flexibility, because abruptly disrupting binaural cues severely impairs sound localization despite intact function in the spared ear ^13–16^. Yet with sufficient experience, humans and animals with monaural hearing loss can relearn to localize sounds, either through adaptive remapping of altered binaural cue–location associations, or by reweighting of spectral cues from the open ear which are typically less informative under binaural conditions ^13–28^. However, the underlying neural mechanisms remain unclear. Indeed, monaural occlusion broadens the spatial receptive fields of individual neurons throughout the auditory pathway, often transforming directionally tuned responses into omnidirectional ones ^29–36^. Consequently, recovery of sound localization is generally assumed to require plasticity mechanisms that adaptively correct the auditory system’s corrupted binaural representations ^28;29;36–38^. Yet a psychophysical paradox complicates this view: Listeners who learn to localize sounds wearing a monaural ear plug typically show little to no “after effect” upon its removal, nor do they continue to rely on monaural cues once binaural hearing is restored ^15;21;38;39^. These data argue against a simple overwriting of prior spatial representations via adaptive plasticity. Rather, the auditory system must maintain access to multiple, non-interfering representations of space that can be flexibly engaged depending on cue availability, akin to context-dependent memories. How such parallel representations might be implemented at the level of neural populations remains unknown.

We addressed this knowledge gap by studying neural population activity in the higher-order, non-lemniscal “shell” layers of the inferior colliculus (IC), an auditory midbrain region implicated in spatial hearing and experience-dependent plasticity ^40–49^. Using two-photon Ca^2+^ imaging in passively listening mice, we tracked neuronal responses to directional sounds under binaural listening conditions, and approximately 2 hours after monaural ear plugging. We found three functionally distinct classes of sound-responsive neurons that accounted for nearly all spatial sound activity and were differentially impacted by monaural hearing loss. Surprisingly, population-level coding of sound location was largely impervious to the abrupt and profound loss of binaural cues. Instead, monaural hearing loss reoriented the population code into a subspace orthogonal to the prior binaural dimensions, while preserving the separation of different sound-source locations. The persistence of spatial information after ear plugging was due to a redistribution of spatially informative activity away from contralaterally selective ON-responsive neurons and toward OFF-responsive neurons that retain spatial selectivity after occlusion, together with sound-inhibited neurons that lack classically defined spatial tuning. Thus, the auditory mid-brain flexibly switches between non-overlapping population codes depending on cue availability. Adaptive spatial hearing may therefore arise by switching to an alternative population code, rather than rebuilding a degraded binaural representation.

## 2 Results

We virally expressed a genetically encoded calcium indicator (GCaMP6f or 8s) in the left IC of adult mice and measured neuronal responses using 2-photon Ca^2+^ imaging as awake, head-fixed mice passively listened to free-field sounds (Figure 1A). We mapped auditory receptive fields at sound locations ± 90° across the frontal azimuth using broadband noise bursts delivered from a movable speaker system (4-35 kHz bandwidth, 65 dB SPL, 500 ms duration; see Methods). Throughout the manuscript, positive and negative angles denote speaker positions contralateral and ipsilateral to the imaged IC, respectively, with 0° corresponding to the midline.

**Figure 1:**
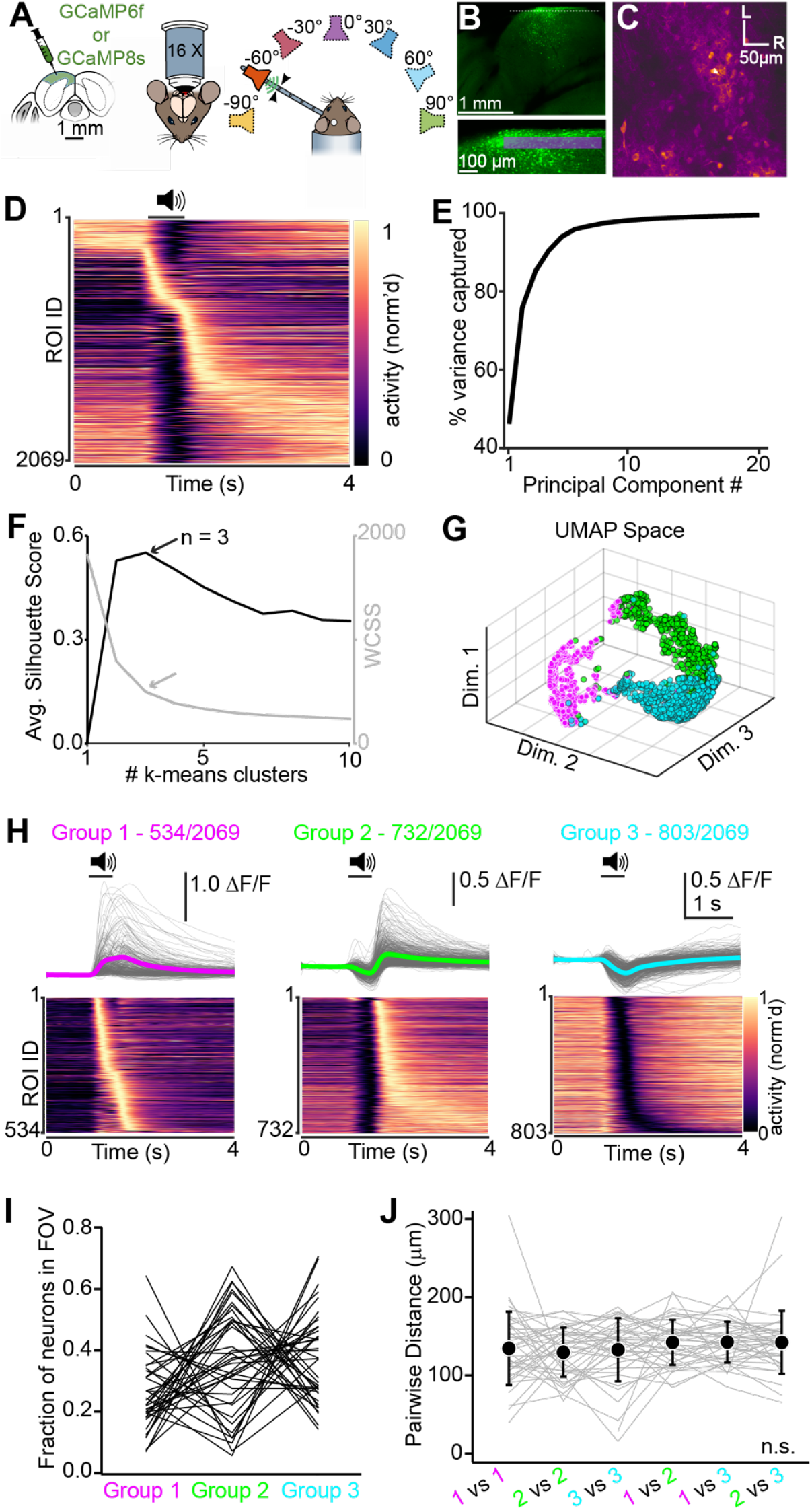
Shell IC activity is dominated by three distinct temporal motifs. A) Cartoon of experiment. Sounds are presented from distinct locations in the frontal azimuth using a movable speaker. B) Example confocal microscopy section of GCaMP8s expression in the shell IC. Lower panel: The area near the dotted line at higher magnification, showing the range of depths for the imaged FOVs across all mice. C) FOV from an exemplar multiphoton imaging session. L = lateral; R = Rostral. D) Heatmap of the normalized fluorescence activity of all sound responsive neurons, averaged across sound locations and ordered according to time of peak. Black bar and loudspeaker denote sound presentation. E) Cumulative variance captured across 20 principal components of the trial-averaged fluorescence time series in (D). F) Silhouette scores (black) and within cluster sum of squares values (WCSS; gray) of the PCA projections plotted as a function of the number of k-means cluster. Arrows, maximum silhouette value and “elbow” of WCSS function indicating k = 3 clusters as optimal. G) Data in D projected into the first three dimensions of UMAP space. Each neuron is color coded according to its group assignment using PCA + k-means. Of note is the separability of the three groups. H) Mean fluorescence (top) and summary heatmap (lower panel) for neurons in Groups 1-3 (left, middle, and right columns respectively). Gray traces are individual neurons. colored traces are group averages. I) Fraction of sound responsive neurons in Group 1, 2, or 3 plotted for n = 41 FOVs. Lines connect individual FOVs. Of note, most FOVs contain neurons in each group. J) Pairwise distances between neurons assigned to same or different groups. Gray lines are individual FOVs, black is mean *±* SEM.

We recorded from n = 3550 regions of interest (ROIs) corresponding to individual IC neurons 30-100 µm from the tectal surface (Figure 1B,C). We quantified neuronal activity as the fluorescence intensity over time (ΔF/F) relative to a 1 s baseline prior to sound presentation (n = 41 fields of view [FOV] in N = 15 mice. Mean ± SEM neurons per FOV: 86.6 ± 5.4. Range: 38-167). An autocorrelation bootstrap analysis identified 2069/3550 (58.3%) of all ROIs as significantly sound responsive under our conditions ^50–53^, and these sound responsive neurons were used for single neuron analyses in Figures 1-3 (Mean ± SEM significant per FOV: 55.8 ± 3.0%. Range: 15.8 – 89.8%).

### 2.1 Three functionally distinct neuron groups account for nearly all shell IC sound responses

As a first pass characterization of shell IC sound responses, we averaged each sound responsive neuron’s fluorescence across all sound locations and ordered the neurons by the time of their peak response. Activity profiles were diverse: Sound presentation either increased or suppressed activity, such that neurons exhibited peak responses before, during or after sound presentation (Figure 1D). This result suggests that IC neuron activity might reflect distinct functional neuron classes defined by their temporal dynamics. To test this hypothesis, we performed principal component analysis (PCA) on the trial-averaged fluorescence time series across neurons. Projection onto the first 20 principal components captured 99.5% of the total variance (Figure 1E), providing a low-dimensional representation of response dynamics. We then applied k-means clustering to these projections, with k = 3 emerging as the optimal clustering solution based on silhouette scores and within-cluster sums of square analyses (Figure 1F). We hereafter refer to these three functional populations as Groups 1-3. Consistent with this classification, UMAP embeddings of the same data revealed clear separation of the three neuron groups discovered by the PCA + k-means pipeline (Figure 1G).

Neurons in each group had strikingly distinct response profiles. Group 1 neurons (534/2069; 25.8%) were excited during sound presentation, with fluorescence typically returning toward baseline upon stimulus offset (Figure 1H, left). This activity pattern is similar to the ON-responsive properties classically described in IC neurons ^54^. In contrast, Group 2 neurons (732/2069; 35.4%) were predominantly suppressed during sound presentation and had a prominent excitatory OFF response (Figure 1H, middle). Group 3 neurons (803/2069; 38.8%) were also suppressed during sound presentation but lacked OFF excitation, thereby resembling the sound-inhibited responses reported in several recent studies of the IC ^50;51;53^ (Figure 1H, right). The three response groups were found in most FOVs (Figure 1I) and intermingled rather than topographically organized, as pairwise soma distances between neurons of the same or different groups were not significantly different (Figure 1J; mixed-effects model, F(3.574,145.8) = 1.478, p = 0.22). Altogether, these results indicate that shell IC activity is parsimoniously explained by three dominant temporal response classes. Interestingly, the Group 1 neurons which resembled ON-responsive neurons often reported in microelectrode studies of the IC ^54;55^ were the minority population in our conditions. Rather, our observations suggest that sound evoked activity in the shell IC relies heavily on inhibition-dominated and temporally structured OFF-response dynamics, rather than classical ON excitation.

### 2.2 Distinct neuron groups have unique spatial tuning properties

Does spatial tuning differ across the three neuron groups? In Group 1 neurons, receptive fields were significantly biased towards the contralateral hemifield (Figure 2A,B; Supplemental Figure 1A. 342/534, or 64.1% with preferred contralateral locations), similar to the spatial tuning properties classically described in IC units ^56–67^. Nevertheless, Group 1 neurons with preferred responses to ipsilateral and midline sound locations were also encountered, indicating that ON excitation in shell IC neurons is not exclusively restricted to the contralateral hemifield (134/534 and 61/534 neurons, respectively; Supplemental Figures 1B,C). In contrast, Group 2 neurons typically had broadly tuned inhibitory ON responses and more spatially restricted excitatory OFF responses (Figure 2C,D). Moreover, in contrast to the contralaterally biased ON excitation of Group 1 neurons, the preferred positions of Group 2 OFF responses were distributed uniformly across the frontal azimuth (Supplemental Figure 2A), with some neurons preferentially responded to intermediate frontal sound locations similar to tuning described in barn owl external IC, brachium, and superior colliculus (Supplemental Figure 2B,C) ^11;68–71^. Finally, most Group 3 neurons were broadly suppressed at all sound locations (Figure 2E,F), though we occasionally observed neurons with clear directional tuning (Supplemental Figure 3) consistent with prior evidence that the IC receives binaurally tuned inhibitory input ^65;72–74^.

**Figure 2:**
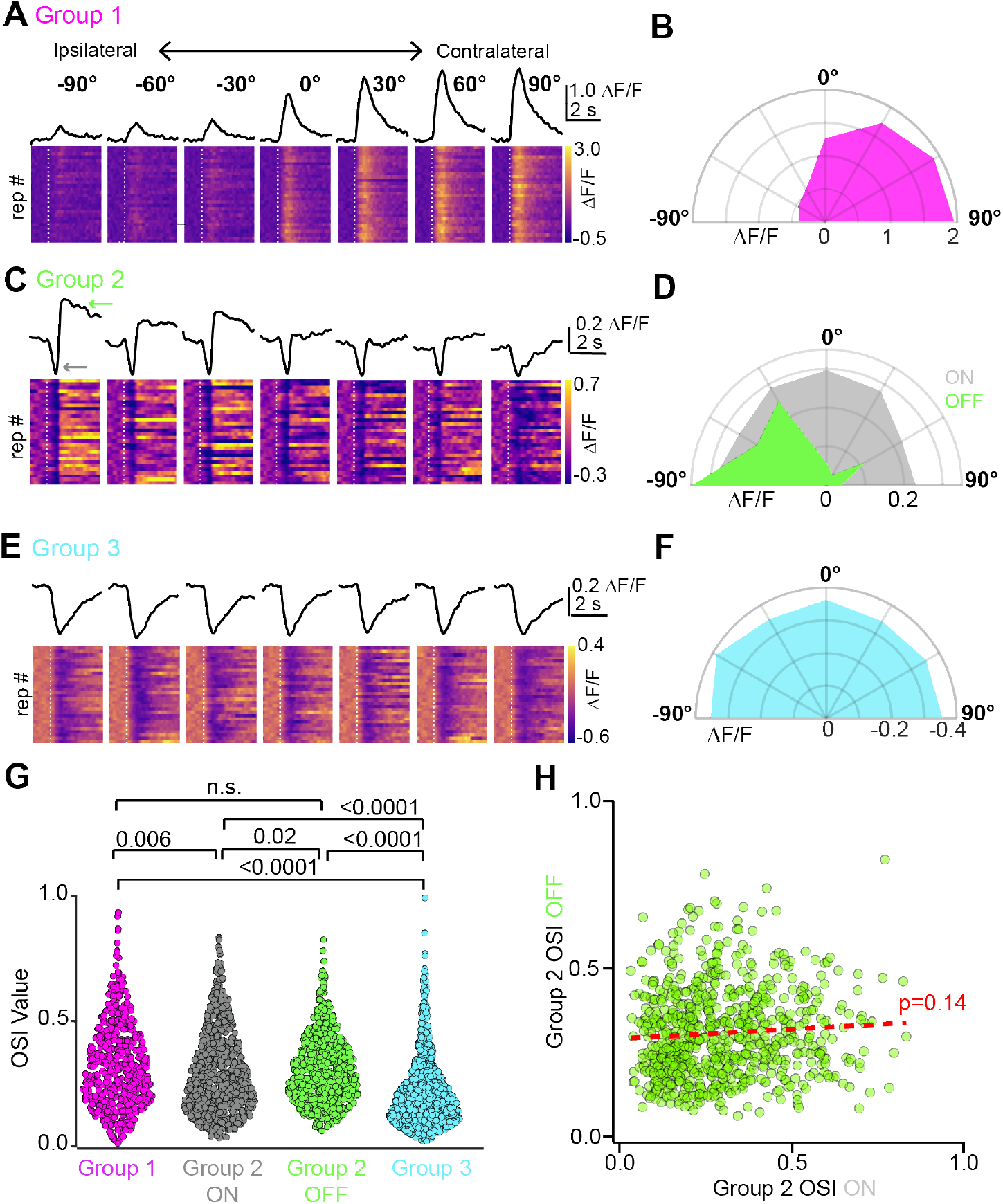
Spatial selectivity differs across Groups 1 - 3. A) Example Group 1 neuron. Black is the mean fluorescence transient, whereas heatmaps show individual trial responses at different sound locations. Dashed line on heatmap is sound onset. B) Half-polar plot of the neuron in (A). The receptive field is primarily restricted to contralateral space. C) Same layout as (A), but for a Group 2 neuron. Gray and green arrows are inhibitory ON and excitatory OFF responses, respectively. Of note, OFF responses are preferentially evoked by ipsilateral sounds. D) Half-polar plot of the ON and OFF receptive field for the neuron in (C). E, F) Same as A-D, but for a Group 3 neuron. G) OSI distributions. Numbers above data clouds are p-values of Bonferroni post-hoc tests. H) OSI values for ON and OFF responses of individual Group 2 neurons are plotted on the X and Y axes, respectively. Of note is the lack of correlation between OSI values (Pearson, r(549) = 0.6271, p = 0.1415, R^2^ = 0.004), indicating that OFF excitation is not a consequence of rebound firing upon cessation of ON inhibition.

**Figure 3:**
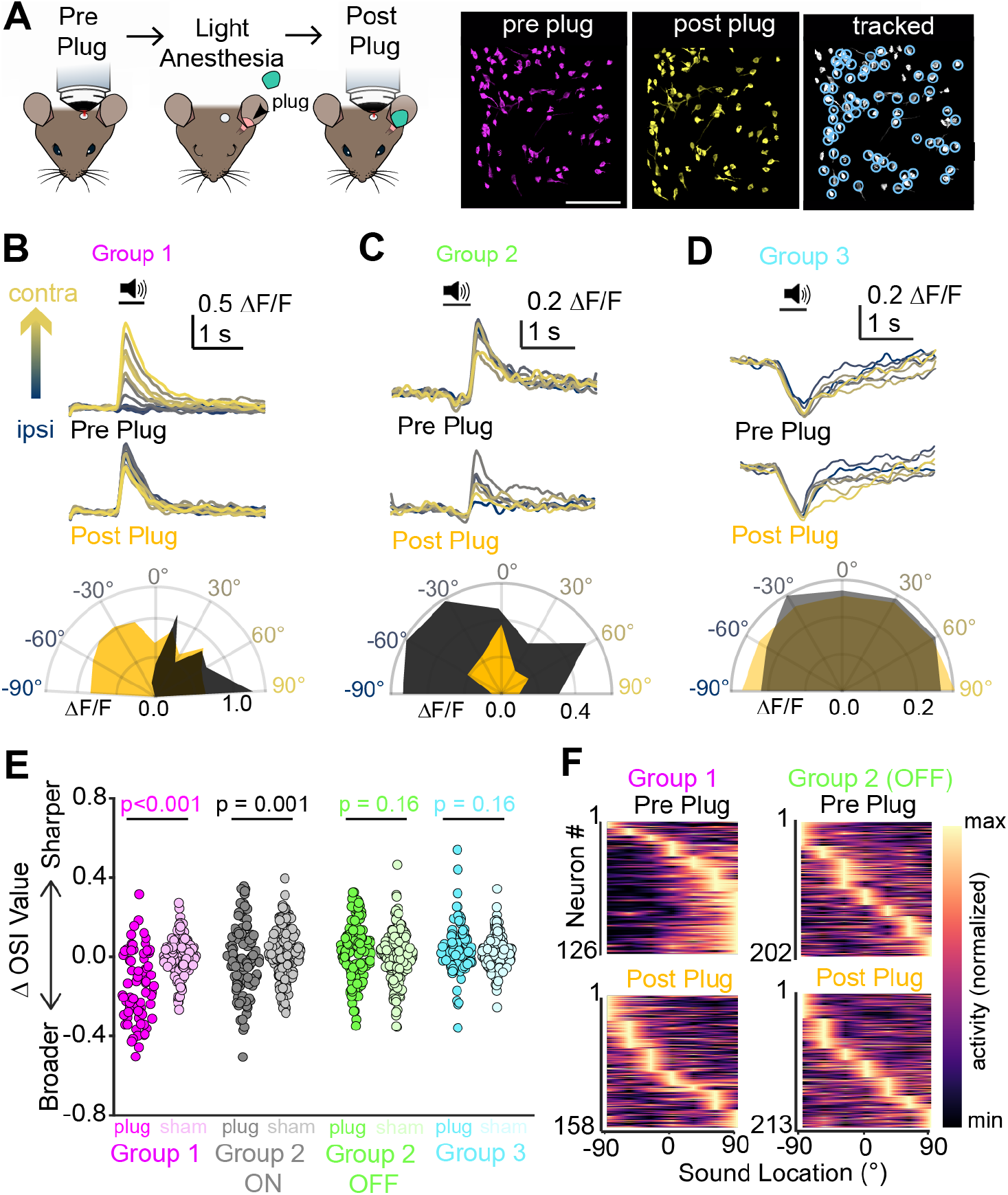
Monaural hearing loss differentially impacts receptive fields of Groups 1-3. A) Left Panel: Cartoon of experiment. Right: Example ROIs in a single FOV tracked across pre and post plug sessions. Scale: 150 µm. B) Upper traces: Example spatial tuning in a Group 1 neuron across pre- and post-plug sessions. Lower: Half-polar plot of the peak DF/F in pre- and post-plug conditions (black and yellow, respectively). Ear plugging unmasks an omnidirectional response in a previously contralaterally tuned neuron. C, D) Same as panel (B) but for example Group 2 and Group 3 neurons. E) Summary data showing DOSI values of single neurons following treatment. P-values above data clouds are post-hoc Wilcoxon rank-sum tests comparing plugged and sham conditions. A two-way ANOVA comparing DOSI values across neuron Group and treatment condition revealed main effects of neuron group (Figure 3H. F(3,936) = 21.31, p<0.001), treatment condition (e.g., ear-plugging vs. sham; F(1,936) = 20.08, p<0.001), and a group x treatment interaction (F(3,936) = 19.19, p<0.001). F) Summary heatmaps of the row normalized excitatory responses at each sound location for all Group 1 (left) and Group 2 neurons (right) in pre and post-plug sessions (upper and lower panels, respectively). Of note, neurons are ordered by preferred position independently for each session.

To further quantify spatial tuning across groups, we computed an Orientation Selectivity Index (OSI) for each neuron ^75^. OSI values range from 0 to 1, reflecting uniform responses across azimuth to responses restricted to a single location, respectively. OSI distributions differed significantly across groups despite substantial within-group variability (Figure 2G. Kruskal-Wallis test, F(3,250), p<0.001. Median OSI values of 0.3, 0.26, 0.29, and 0.18 for Group 1, Group 2 ON, Group 2 OFF, and Group 3, respectively). Excitatory ON responses of Group 1 neurons and OFF responses of Group 2 neurons had comparable OSI value distributions, whereas both excitatory responses were significantly more selective than the inhibitory ON responses of Group 2 and Group 3 neurons (Figure 2G). Inhibitory ON responses in Group 2 neurons were themselves modestly but significantly more selective than those of Group 3 neurons (Figure 2G), suggesting that shell IC neurons receive multiple inhibitory inputs with distinct spatial properties rather than a single uniform source of broadly suppressive drive. Importantly, Group 2 ON and OFF selectivity were not significantly correlated (Figure 2H), arguing against the idea that OFF excitation simply reflects rebound firing following spatially tuned inhibition. Rather, the data suggest that ON and OFF responses arise from distinct presynaptic inputs converging onto the same neuron ^76–78^. Consistent with this interpretation, a minor subset of Group 1 and Group 2 neurons exhibited both ON and OFF excitation (Supplemental Figure 4A,B; n =108 neurons found in N = 28 FOVs across N = 13 mice. Fraction Group 1: 73/108, 67.6%); the paucity of this response type likely explains the limits of our clustering method’s ability to adequately separate these neurons from the other sound-excited populations. Within these neurons, preferred ON and OFF sound locations were frequently separated by more than 60° and differed significantly from a hypothetical median of zero (Supplemental Figure 4C, one sample Wilcoxon test, z = 8.36, p < 0.001), further supporting the idea that ON and OFF excitation originate from distinct presynaptic inputs. Altogether, these analyses show that at the single neuron level, spatial selectivity appears primarily driven by excitatory responses, regardless of whether said excitation occurs during sound presentation or upon sound offset. However, the preferred sound locations of ON and OFF responses differentially emphasize contralateral versus full-field representations.

**Figure 4:**
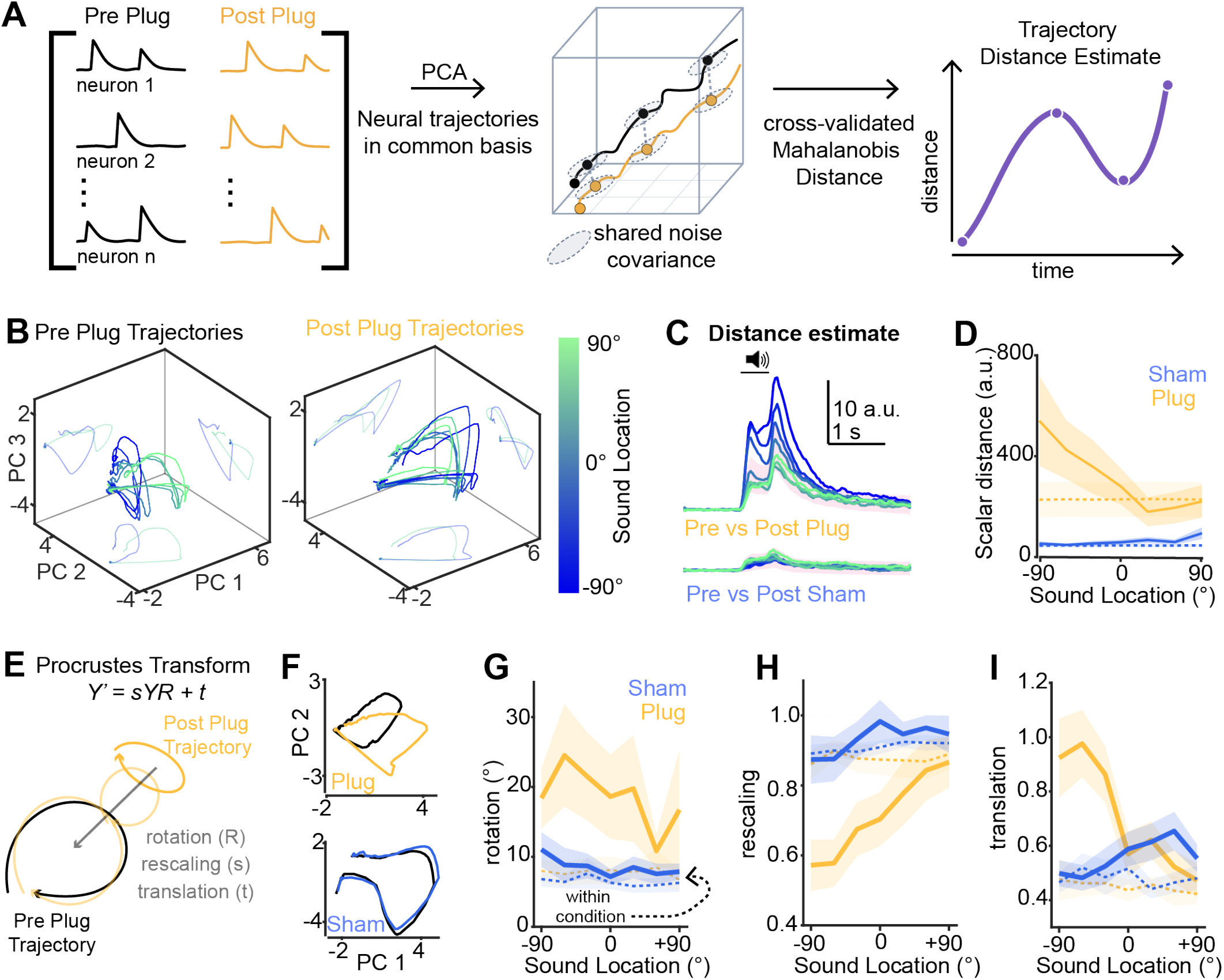
Monaural hearing loss rotates neural trajectories. A) Cartoon of analysis pipeline for panels B-D. B) Example neural trajectories in 3D PCA space (single mouse). “Wall shadows” are the equivalent data on 2D axes. The means of ipsi- and contra-lateral positions are plotted for clarity. C) Cross-validated Mahalanobis (“crossnobis”) distances between pre- and post-treatment trajectories for each sound location in ear-plugged and sham-treated mice (upper and lower traces, respectively). Pink shading shows boundaries of crossnobis distances calculated from shuffled null models. Of note, the largest distances are for pre- and post-plug ipsilateral positions, whereas the distance between contralateral positions is similar to a shuffled null model. D) Summary plot of crossnobis distances at each sound location for ear-plugged and sham-treated mice (yellow and blue, respectively). Shading is ± SEM, dotted lines are null shuffles. E) Cartoon of Procrustes analysis. F) Representative 2D neural trajectories from plug- and sham mice before and after treatment. Trajectories are averaged across all sound locations for clarity. G) The rotation term of the Procrustes transformation plotted per sound location for plug-(yellow) and sham-treated (blue) mice. Solid lines are mean transformations required to align pre- and post-treatment trajectories. Shading is ± SEM. Dotted lines are within-condition controls in which trajectories were generated from separate halves of the data and aligned within the same condition (30 repeats). H, I) Same as (G), but for the rescaling (H) and translation (I) variables.

### 2.3 Monaural ear plugging differentially impacts spatial tuning of Groups 1-3

Ipsilateral sounds typically suppress IC neuron spiking to sounds presented to the dominant, contralateral ear ^65;72;79–82^; this phenomenon is thought to contribute to interaural level difference (ILD) coding and the contralateral bias of spatial receptive fields ^83–86^. Given the striking diversity of spatial tuning under binaural conditions, we asked whether monaural ear-plugging would equally, or differentially, impact single neuron spatial tuning across groups.

To test this idea, we measured the spatial receptive fields of shell IC neurons before, and within ∼2 hours after inserting an ipsileratal ear plug (Figure 3A). This manipulation caused an average threshold shift of ∼45-55 dB SPL at the plugged ear across frequencies tested via pure-tone auditory brainstem response measurements (N = 3 mice). Individual neurons were tracked across pre- and post-plug sessions using an offline registration approach ^87^. Within-cell comparisons of receptive field properties were quantified as DOSI values relative to the pre-plug baseline, with the ON and OFF receptive fields of Group 2 neurons measured separately; Positive and negative DOSI values reflect sharpening and broadening of receptive fields, respectively. To control for non-specific effects, we compared ear-plug experiments (n = 10 FOVs from N = 10 mice) with sham-treated controls (n = 12 FOVs from N = 8 mice).

Ear-plugging differentially impacted single neuron tuning selectivity. In Group 1 neurons, ear plugging typically broadened receptive fields compared to sham treatment, with contralateral receptive fields often becoming omni-directional (Figure 3B,E. Median DOSI values: -0.188 vs. 0.004, p < 0.001 Wilcoxon ranksum test. n = 75 and 119 neurons from ear-plug and sham mouse groups, respectively). This result is consistent with the classic interpretation that contralateral azimuth tuning arises from “EI” interactions, with ipsilateral sounds constraining spiking to the contralateral hemifield ^34;65;72;83;84;88–91^. Markedly different effects occurred in Group 2 neurons (Figure 3C,E). DOSI values of inhibitory ON responses were modestly, though significantly lower in ear-plugged compared to sham treated mice (Median DOSI values: -0.027 vs. 0.045, p = 0.001, Wilcoxon rank-sum test. n = 112 and 146 neurons from ear-plug and sham mice, respectively). By contrast, excitatory OFF responses showed substantial heterogeneity in the magnitude and direction of change across individual cells. Some Group 2 neurons had receptive fields which sharpened after ear-plugging (Figure 3C; Supplemental Figure 5A), remained broadly tuned (Supplemental Figure 5B), or shifted their preferred location (Supplemental Figure 5C). Thus unlike in Group 1 neurons, ear-plugging did not systematically decrease the OSI values of Group 2 neuron OFF responses (median DOSI values: 0.014 vs -0.005, p = 0.16 Wilcoxon rank sum test. n = 112 and 146 neurons from ear-plug and sham-treated mice, respectively). Rather, OFF on average maintained spatial selectivity following an abrupt and profound reduction in binaural cues. Additionally, the broadly suppressive inhibition of Group 3 neurons remained stable after ear plugging (Figure 3D,E. Median DOSI 0.029 vs. 0.012, p = 0.16 Wilcoxon rank-sum test, n = 90 and 144 neurons from ear-plug and sham-treated mice, respectively). This result is consistent with the relatively weak spatial selectivity of inhibitory responses in our conditions. Collectively, the results show that rather than uniformly degrading shell IC spatial tuning, monaural hearing loss selectively reduces selectivity and abolishes the contralateral preference of Group 1 neurons preferred locations. By contrast the distribution of preferred OFF-response locations, which span the frontal azimuth, appeared preserved across the Group 2 neuron population (Figure 3F).

**Figure 5:**
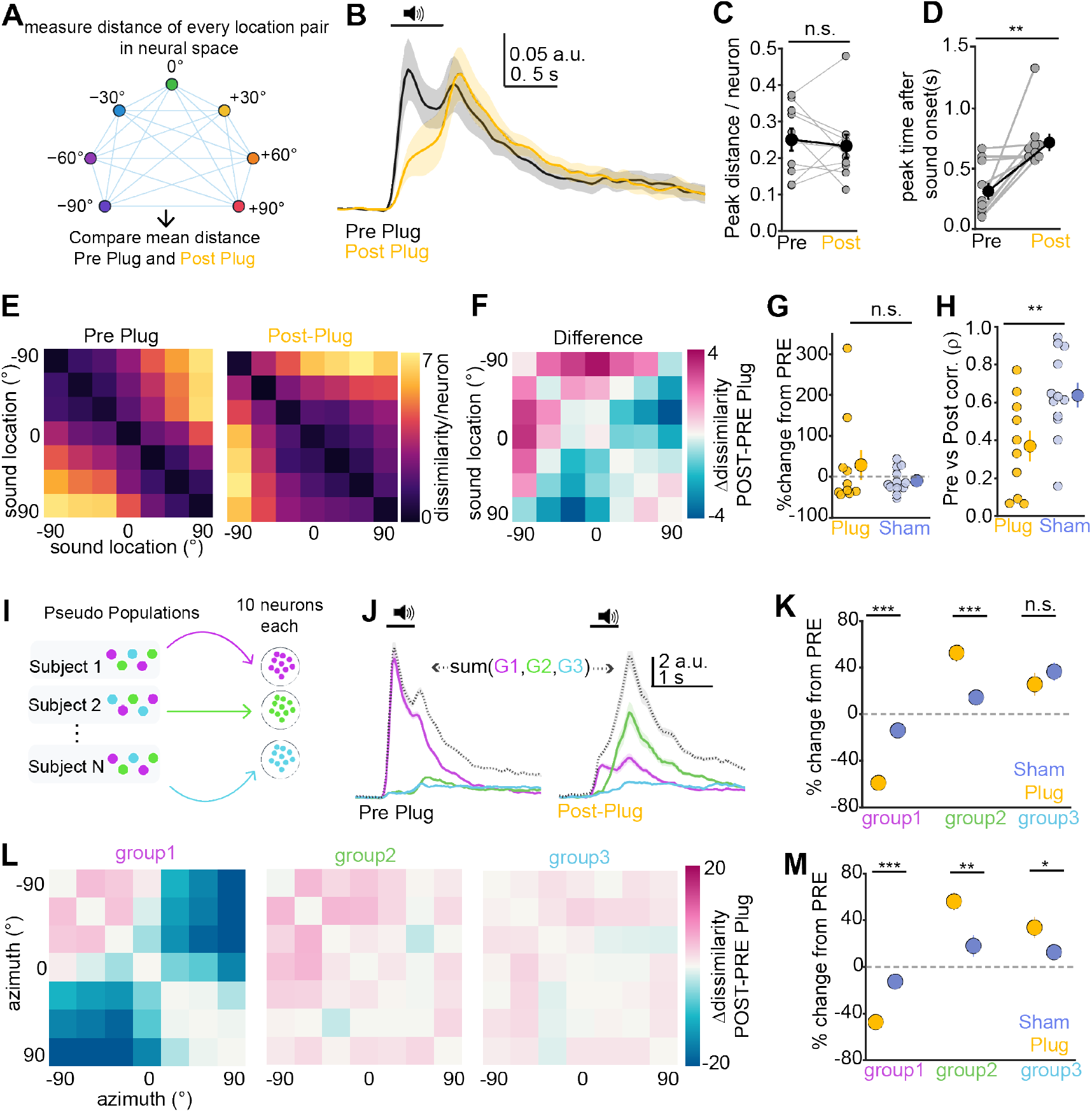
Distinct neuron groups contribute to trajectory separation before and after ear plugging. A) Cartoon of analysis. B) Mean neural distance over time, pre- and post-plug (black and yellow, respectively). Shading is ± SEM. C) Summary of peak distance. Gray is individual experiments, black is mean ± SEM. D) Summary, time of peak following sound onset in pre- and post-plug conditions. Colors are same as in (C). E) RDMs for pre- and post-plug conditions, averaged across all ear-plug experiments. F) Difference of pre- and post-plug RDMs. Warm and cool colors are sound location pairs that become more or less dissimilar after ear-plugging, respectively. G) Summary, mean % change in non-diagonal RDM pairwise dissimilarity relative to pre-treatment, for ear-plug and sham conditions (yellow and blue, respectively). Small symbols are individual mice, large symbols are mean ± SEM. H) Pearson’s r correlation of pre- and post-RDMs. Symbols and colors are same as (G). I) Pseudo-populations are generated by sampling without replacement equal counts of Group 1, 2, and 3 neurons. J) Mean distances for pseudo-populations of Group 1, 2, and 3 neurons, in pre- and post-plug conditions (left and right, respectively). Dotted lines are the linear sum across the three groups. Shading is 95% confidence interval (C I). K) Mean %change in peak distance relative to pre-treatment condition for pseudo-populations drawn from ear-plugged or sham-treated experiments (yellow and blue, respectively). L) RDM difference matrices for Group 1-3 pseudo-populations of ear-plugged mice. M) % change across RDM entries relative to pre-treatment. Colors and symbols are same as in (K).

### 2.4 Neural population trajectories remain separable after monaural hearing loss

The heterogeneous single-neuron effects described above raise a key question: does monaural deprivation degrade spatial coding at the population level, or does spatial information persist despite reorganization of ensemble activity? To address this, we measured IC population activity as low-dimensional neural trajectories in PCA space, and quantified how these trajectories changed after ear plugging. Neural trajectories provide a compact description of time-varying ensemble activity by projecting high-dimensional population responses onto a smaller set of basis patterns; this allows the geometry and dynamics of ensemble activity to be compared across sound locations and treatment conditions in a common, low-dimensional space ^92–94^. For these analyses, we included all neurons that could be reliably tracked across pre- and post-treatment sessions, regardless of whether individual neurons met our *a priori* criteria for sound responsiveness. Effective dimensionality of the datasets was comparable across pre- and post-treatment conditions, and across plug and sham cohorts (Supplemental Figure 6). Acute monaural hearing loss therefore does not change the overt dimensionality of sound-driven population activity in our paradigm.

**Figure 6:**
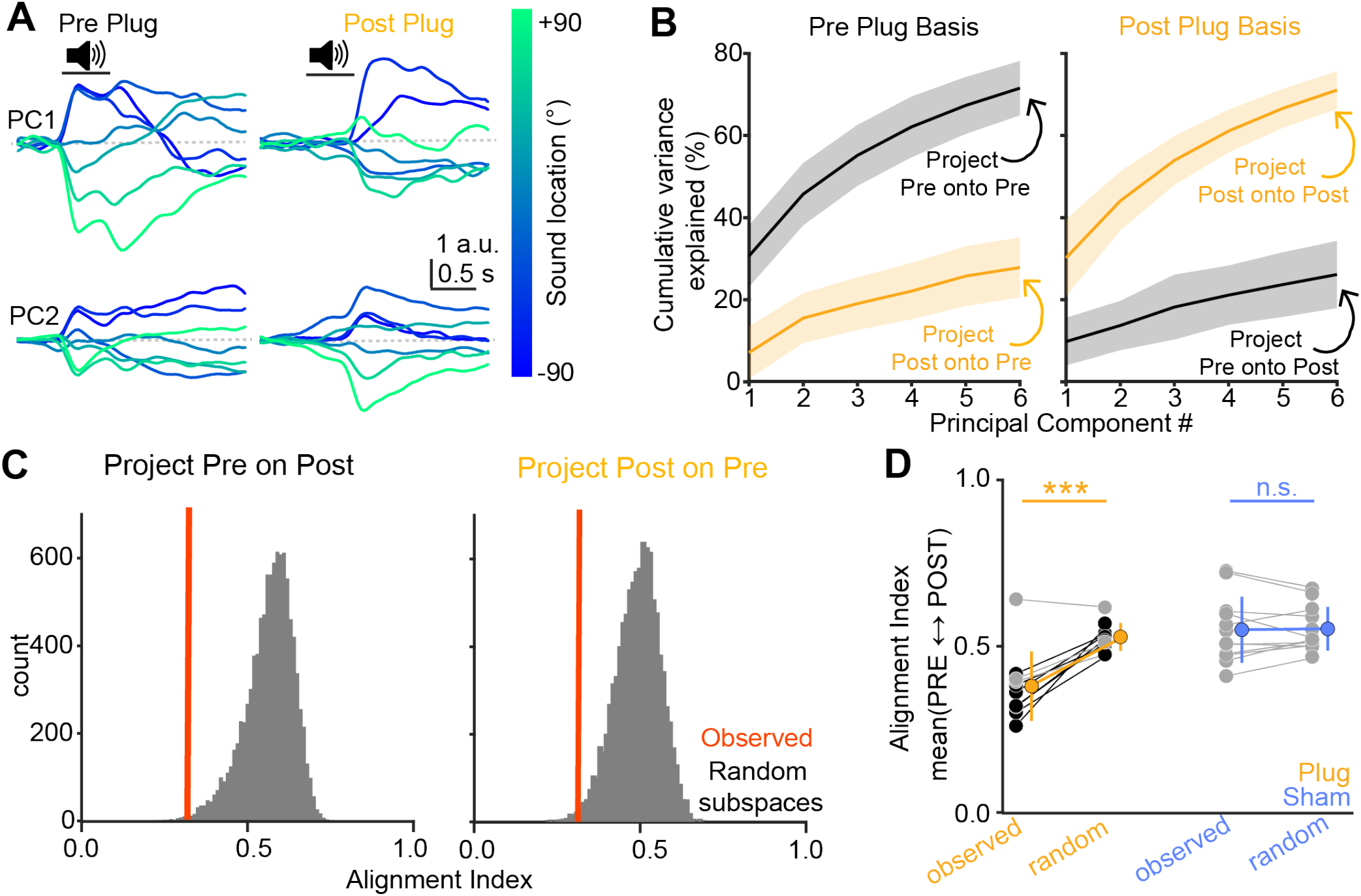
Monaural hearing loss reorients population coding subspaces. A) Separate bases were fit to the pre- and post-plug activity matrices, and population activity was projected onto the first two PCs. Data are from a single subject. B) Left: Cumulative variance explained when pre- and post-plug activity is projected onto the basis discovered from pre-plug data (black and orange, respectively). Right, same, but when pre- and post-plug activity is projected onto the post-plug basis. Lines are group means and shading indicates ± SD across 10 experiments. C) Alignment indices for the example experiment in (A), calculated by projecting pre-plug activity onto the post-plug subspace (left) and post-plug activity onto the pre-plug subspace (right). Vertical red lines are the observed alignment index. Gray histograms is distributions generated from 10,000 random subspaces drawn under a covariance-preserving null model, following Elsayed et al. (2016). Lower alignment indices indicate less overlap. D) Summary of observed and random alignment indices in plug and sham experiments (yellow and blue, respectively). Large symbols and error bars show mean ± SD. Connected points are individual experiments. Black symbols are individual experiments in which the observed alignment index differed significantly from the random-subspace distribution; gray symbols are nonsignificant experiments.

Different sound locations generated distinct trajectories under binaural conditions, with ipsilateral and contralateral locations occupying separable regions of the low-dimensional population state space (Figure 4B). Following ear plugging, however, trajectories evoked by ipsilateral sounds shifted markedly, encroaching on regions typically occupied by contralateral positions, while contralateral trajectories appeared comparatively less altered (Figure 4B). We quantified this reorganization by calculating the cross-validated Mahalanobis distance (crossnobis) ^95–97^ between pre- and post-plug trajectories at each sound location, and comparing the crossnobis timecourses to sham-treated controls. Following ear plugging, pre–post distances increased sharply during sound presentation and exhibited a second increase upon sound offset, indicating that both ON and OFF population dynamics were altered by earplugging (Figure 4C). Pre–post distances were substantially greater in ear-plugged compared to sham-treated mice, with the largest differences appearing at sound locations nearest to the ipsilateral, plugged ear. In comparison, pre–post changes in sham-treated mice showed minimal location dependence and were similar to distances estimated from shuffled null data (Figure 4C). We quantified these results by summing the crossnobis distance during the sound ON and OFF period at each sound location, and comparing the values between ear-plugged and sham treatment. A two-way ANOVA revealed significant main effects of sound location (F(6,120) = 3.86, p = 0.044) and treatment condition (F(1,120) = 11.88, p = 0.003), and a sound location × treatment interaction (F(6,120) = 5.05, p = 0.020). Thus, ear-plugging differentially alters neural trajectories corresponding to distinct sound locations, with the largest population-level reorganization occurring for sound locations near the plugged, ipsilateral ear.

We further quantified how ear-plugging differentially alters ipsi- and contra-lateral neural trajectories using Procrustes analysis, which determines the transformation that optimally aligns two dimensional trajectories by decomposing their geometric relationship into rotation, uniform rescaling, and translation variables (Figure 4E). Consistent with the population trajectories visualized above, substantial transformations were required to align pre- and post-plug trajectories, whereas pre- and post-sham trajectories were more similar (Figure 4F). We separately compared each component of the Procrustes transformation across sound location and treatment condition using two-way mixed-design ANOVAs (Figures 4G–I). For the rotation variable (Figure 4G), we found a significant main effect of treatment condition (F(1,20)=4.53, p = 0.046), no main effect of sound location (F(6,120)=1.33, p = 0.28), and no interaction effect (F(6,120) = 0.92, p = 0.41). Thus, monaural hearing loss rotates the orientation of sound-evoked neural trajectories, though the absolute magnitude of this rotation is independent of sound location. By contrast, the scaling variable (Figure 4H) showed significant main effects of treatment condition (F(1,20) = 7.61, p = 0.012), sound location (F(6,120) = 2.8, p = 0.03), and a treatment condition x sound location interaction (F(6,120) = 7.91, p <0.001). Similar results were observed for the translation variable (Figure 4I. Treatment condition: F(1,20) = 4.48, p = 0.047; sound location (F(6,120) = 3.07, p = 0.021; treatment condition x sound location interaction: F(6,120) = 8.16, p <0.001). Scaling and translation effects were greatest at ipsilateral sound locations, suggesting that location-dependent changes in trajectory distance and shape contribute strongly to the hemifield specific differences between pre- and post-plug trajectories (Figure 4C,D). As a control to estimate the transformation expected from finite trial sampling, we also computed Procrustes alignment between trajectories independently generated from separate halves of the data within each experimental condition (Figure 4G–I, dotted lines). Two-way mixed-design ANOVAs revealed no significant main effects of sound location or treatment condition for any of the Procrustes variables calculated from these within-condition transformations (all *p* > 0.05). Thus, the treatment- and location-dependent changes observed for the pre–post transformations cannot be accounted for by sampling variability alone. Instead, monaural hear loss differentially reorganizes several features of population trajectories: The changes in trajectory scale and position are strongest for sounds in the hemifield near the occluded ear, whereas trajectory orientation change with a more uniform magnitude across the frontal azimuth.

### 2.5 Monaural hearing loss redistributes spatial information across time and neuron groups

Do ear-plug related changes in neural trajectories reduce spatial discriminability, or can neural populations nevertheless distinctly represent sound locations after acute degradation of binaural cues? To address this question, we quantified the distance in full neural space between population trajectories, for all 21 pairwise combinations of the seven sound locations. We then compared the time-resolved, mean trajectory separation across pairs before and after ear-plugging (Figure 5A). Before ear plugging, trajectory separation increased rapidly following sound onset, declined during the sustained response, and exhibited a smaller secondary increase following sound offset (Figure 5B). By contrast after ear-plugging, the initial separation rise was markedly slowed and a prominent second phase emerged around sound offset. Consequently, average peak separation between sound location trajectories was comparable before and after plugging (Figure 5C. Mean ± SEM 0.25 ± 0.03 vs 0.23 ± 0.03 for pre and post-plug. n = 10 FOVs from N = 10 mice, p = 0.55, sign-rank test). However, the latency of maximal separation significantly shifted later after sound onset in post-compared to pre-plug sessions (Figure 5D; Mean ± SEM 0.33 ± 0.07 vs 0.72 ± 0.08 ms for pre and post-plug sessions, p = 0.004, sign-rank test). Importantly the shift in latency results was not observed in sham treated mice (data not shown; 0.34 ± 0.08 vs 0.48 ± 0.08 ms. N = 12 FOVs from N = 8 mice, p = 0.12, sign-rank test): Thus, despite a profound reduction in binaural cue availability, IC populations can nevertheless generate location-specific population activity patterns, albeit with a peak discriminability shifted towards OFF-dominated population activity.

Does comparable trajectory separation across hearing conditions reflect a preserved distance relationships between sound locations, or does monaural hearing loss cause a redistribution separability across location pairs? To address this question, we constructed representational dissimilarity matrices (RDMs), in which the values of each off-diagonal matrix element quantifies the dissimilarity between population activity evoked by a particular pair of experimental conditions ^98^. This approach summarizes the geometric relationship of the neural representations between different sound locations, whereby larger values indicate more distinct population responses between pairs of sound locations. Under binaural conditions, RDMs had a structured organization whereby dissimilarity increased as a function angular separation, with particularly strong dissimilarity across the two hemifields (Figure 5E). This graded organization changed after ear-plugging: High dissimilarity remained between ipsilateral pairs, whereas ipsilateral–contralateral pairs exhibited lower dissimilarity values. Consequently, difference matrices between pre- and post-plug RDMs revealed both increases and decreases in pairwise dissimilarity rather than a uniform collapse of spatial separation (Figure 5F), and the average percentage change in pairwise dissimilarity was not significantly different between plug and sham cohorts (Figure 5G; 29.8 ± 37.8% vs -11.01 ± 8.1% for plug and sham conditions, respectively. P = 0.97, rank-sum test). However we observed a marked reduction in the correlation between pre- and post-treatment RDMs in plugged compared to sham mice (Figure 5H. Mean ± SEM Pearson’s r of pre- and post-treatment RDMs: 0.36 ± 0.08 vs 0.7 ± 0.08, for sham and plug conditions p = 0.008, unpaired t-test). Monaural hearing loss therefore preserved the overall magnitude of dissimilarity across sound locations while nevertheless reorganizing the internal structure of a prior binaural population code.

The opposing changes in pairwise separation between sound location pairs could arise from the differential single cell effects of ear plugging seen in the three distinct neuron groups. We tested this hypothesis by constructing pseudo-populations, sub-sampled across subjects and containing equal numbers of Group 1, 2, and 3 neurons that could be reliably tracked across pre- and post-plug sessions (Figure 5I). We then calculated distances between all sound-location pairs, and decomposed the population distance into contributions from each group. This approach controlled the number of sampled neurons when measuring time-resolved distances and comparing across neuron groups and treatment conditions. Before ear-plugging, Group 1 neurons made the largest contribution to population separation, whereas Groups 2 and 3 contributed smaller components (Figure 5J). Ear-plugging inverted this relationship, such that Group 1 neurons’ contribution was strongly reduced, while Group 2 trajectory distances increased and shifted toward the sound-OFF period; Group 3 exhibited a smaller increase (Figure 5J). Relative to sham treatment, ear-plugging significantly reduced the peak distance of Group 1 pseudo-populations from the pre-plug baseline, while significantly increasing peak distances of Group 2 pseudo-populations; peak distances of group 3 neurons did not change significantly after plugging compared to sham treated mice (Figure 5K. Mean ± SEM % change relative to pre-treatment for plug vs sham: Group 1, -59.0 ± 4.2 vs -13.9 ± 3.8, p<0.001. Group 2, 52.8 ± 7.4 vs 14.4 ± 6.9, p<0.001. Group 3, 25.7 ± 9.8 vs 36.6 ± 7.8, p = 0.39. Unpaired Welch t-tests). Sound location-specific neural trajectory distances therefore shifted away from the ON-dominated Group 1 neurons under binaural conditions, towards OFF-responsive Group 2 neurons following ear-plugging.

Group-specific RDMs provided a complementary view across individual sound-location pairs. Group 1 RDMs exhibited both local increases and decreases in dissimilarity and an overall reduction in pairwise separation after ear plugging (Figure 5L). In contrast, pairwise separation increased substantially within Group 2, and more modestly within Group 3 (Figure 5L,M. Mean ± SEM % difference in off-diagonal entries for plug vs sham conditions. Group 1: -47.2 ± 5.4 vs. 12.5 ± 3.9, p<0.001. Group 2: 56.2 ± 7.5 vs 18.0 ± 9.3, p=0.004. Group 3: 33.8 ± 9.0 vs 12.7 ± 4.6, p = 0.04, unpaired Welch t-tests). These results argue that the preserved population separation after monaural hearing loss emerges from opposing changes across functional neuron groups, rather than a uniform maintenance of the pre-plug representation. Consequently, maximal population separation shifts toward sound offset, and the pairwise geometry is reconfigured in a manner that maintains a similar mean representational separation between sound locations as in binaural conditions. Thus, abrupt monaural hearing loss rapidly rebalances which neurons functionally contribute to sound location coding in the IC.

### 2.6 Monaural hearing loss re-orients population coding subspaces

Does the geometric organization of population activity after monaural hearing loss occurred within the same population dimensions engaged under binaural hearing conditions, or does it instead reflected a shift toward distinct neural subspaces? To address this question, we independently identified the dominant population dimensions in pre- and post-plug sessions via PCA, and quantified the extent to which population activity from each hearing condition aligned with the subspace identified in the other condition.

Population activity followed visibly different low-dimensional trajectories before and after ear plugging: PCs 1 + 2 diverged rapidly following sound onset in pre-plug conditions, whereas in post-plug conditions PCs showed markedly weaker early separation and separated maximally upon sound offset (Figure 6A). Moreover, the leading pre-plug dimensions captured substantially less variance of post-plug activity compared to pre-plug activity, and the converse was similarly true for the post-plug dimensions: pre-plug variance was poorly captured by the population dimensions that dominated after plugging (Figure 6B). A two-way repeated-measures ANOVA comparing variance explained across the first six PCs for within-basis projections (PRE-in-PRE vs POST-in-POST) and across-basis projections (PRE-in-POST vs POST-in-PRE) revealed main effects of projection type (F(1,9) = 263.7, p < 0.001), PC number (F(5,45) = 91.6, p < 0.001), and a projection type × PC number interaction (F(5,45) = 42.5, p < 0.001). Thus, the dimensions carrying spatially structured activity change substantially following monaural hearing loss. We further quantified this re-organization using the subspace alignment index ^93;99^, which measures how much of the dominant variance in one population response can be captured by a sub-space derived from another response. An alignment index of 1 indicates complete correspondence between the relevant population dimensions, whereas progressively lower values indicate increasingly distinct sub-spaces. We then compared the observed alignment to distributions obtained from 10,000 random subspaces sampled according to the covariance structure of the combined population activity (Figure 6C; see also ref 95). Alignment indices were significantly lower than their covariance-matched random distributions for at least one cross-projection direction in 7/10 and 0/12 individual ear-plugging and sham control experiments, respectively (Figure 6D; see also methods). A two-way ANOVA revealed a main effect of treatment condition (F(1,20) = 9.17, p = 0.007), alignment type (observed versus random, F(1,20) = 26.05, p<0.001) and a treatment x alignment type interaction (F(1,20) = 23.99, p<0.001). Pairwise post-hoc tests indicated that observed values were significantly lower than the random model for ear-plugged, but not sham-treated mice (Figure 6D. Mean ± SEM for plug and sham cohorts: 0.38 ± 0.03 vs 0.53 ± 0.01; 0.55 ± 0.03 vs 0.55 ± 0.02. p<0.001 and p = 0.82„respectively, post-hoc paired t-test). Thus, monaural hearing loss reorganizes the population code into a different configuration of neural dimensions rather than redistributing activity within the existing binaural subspace.

### 2.7 Monaural hearing loss rotates the binaural coding axis

Are neural representations from ear-plugged mice sufficient to encode sound location? If so, to what extent does binaural and monaural representations rely on overlapping neural features? To test these ideas, we asked if binary classifiers could determine if a sound originated in the ipsi- or contra-lateral hemifield using single trial neural population activity, and whether such classifiers could generalize across pre- and post-plug hearing conditions (Figure 7A). Although a prior study suggested that the high dimensionality of shell IC neural population activity prevents linear classifiers from accurately decoding sound location ^100^, support vector machines (SVM) trained on single trial fluorescence data from pre-plug FOVs performed with high accuracy when tested within condition on held out pre-plug trials (Figure 7B). Additionally, pre-plug classifiers showed little bias when classifying midline trials that were held out of the training fold (Supplemental Figure 7A). In contrast, classifiers trained on pre-plug activity generalized poorly when tested on post-plug trials, and performance fell to near chance similar to shuffled classifiers (Figure 7B. Mean accuracy 81.3% versus 53.5%; 95% confidence intervals [CI] 70.4-89.5% versus 48.8-58.4% for train-pre/test-pre and train-pre/test-post, respectively; n = 10 FOVs from N = 10 mice). Similarly weak cross-condition generalization was observed when classifiers were trained on post-plug trials and tested on pre-plug trials (mean: 58.3%, 95% CI: 52.8-64.9%). This near-chance cross-condition performance was accompanied by a strong contralateral bias (Supplemental Figure 7A), paralleling the open-ear bias observed in humans after acute monaural ear-plugging ^18^. The poor cross-condition decoding did not, however, reflect a loss of sound location information after plugging. Classifiers trained and tested on post-plug trials performed well above chance (mean: 74.0% 95% CI: 67.9-79.5%), showed no systematic hemifield bias (Supplemental Figure 7A), and exhibited “psychometric” slopes comparable to classifiers trained and tested on pre-plug data (Supplemental Figure 7B). Importantly, the loss of cross-condition generalization could not be explained by nonspecific effects of anesthesia, repeated imaging, or representational drift, because classifiers readily generalized across sessions in sham-treated mice (Figure 7B; n = 12 FOVs from N = 8 mice). A two-way, mixed-design ANOVA comparing ear-plugged and sham-treated groups revealed main effects of treatment group (sham vs. plugged, F(1,20)=10.69, p=0.0038), classifier train + test condition (F(3,60)=30.69, p<0.001), and significant treatment group x classifier condition interaction (F(3,60)=13.31, p<0.001), with only the cross-generalization conditions showing significant post-hoc differences between sham and plug treatment (p<0.01, unpaired t-test).

**Figure 7:**
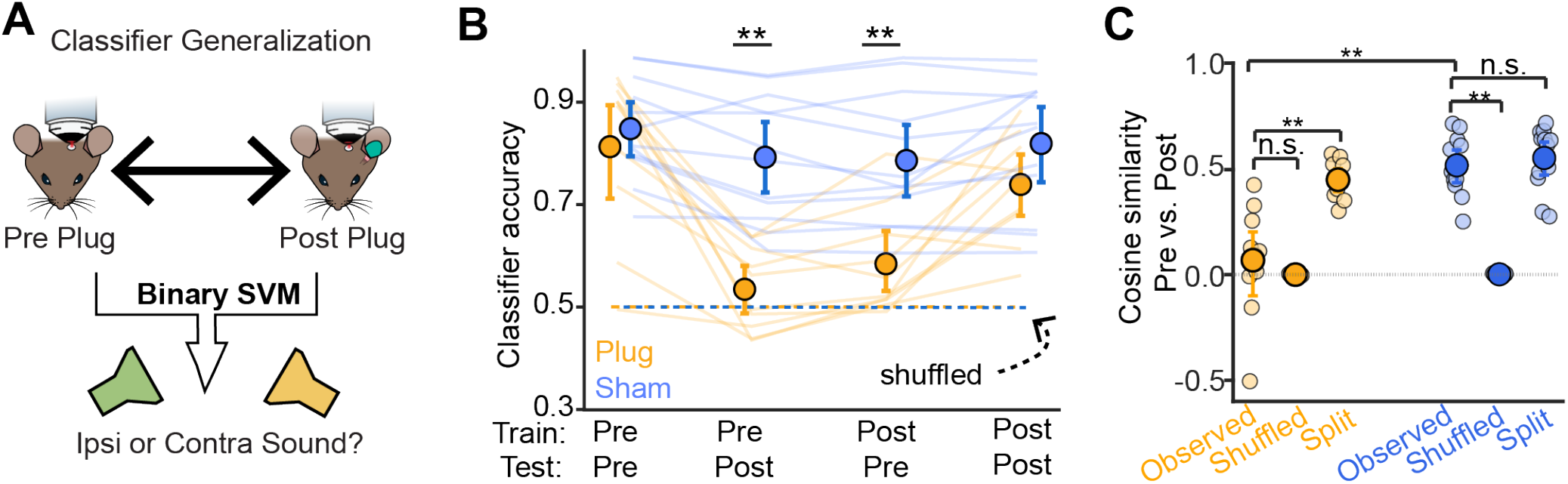
Orthogonal coding axes of azimuthal space under binaural and monaural conditions. A) Cartoon of analysis. Binary SVMs are trained to classify sound location hemifield using neural population activity in pre- or post-plug neuron sessions, and tested for cross-generalization across hearing conditions. B) Summary SVM performance per training and testing condition. Error bars are ± 95% confidence intervals from 1000 bootstrapped iterations of the subject mean. Dotted lines are means of classifiers trained on shuffled class labels. Asterisks: p<0.01, Fisher’s least significant difference post-hoc tests between sham and plug conditions. C) Cosine similarity indices for the binary decision vectors of SVMs trained on pre- and post-manipulation sessions. Ear-plugged and sham-treated mice are in yellow and blue, respectively. Also shown are values of shuffled-null controls and within-condition splits, i.e. the cosine similarity of decision vectors for classifiers trained on separate halves of the data from the same condition. Split values are averaged across pre- and post-conditions. Small points are individual subjects, large points and lines are mean ± 95% confidence interval. Asterisks denote statistical significance of post-hoc tests at p<0.01, n.s. not significant.

The results above indicate that the IC activity after monaural hearing loss is sufficient for a downstream readout to decode sound source hemifield, though the monaural and binaural representations may be poorly aligned. To test this hypothesis, we asked whether ear-plugging rotated the population axis separating ipsilateral from contralateral space. Because the SVM weight vector defines the direction in neural activity space that best separates the two hemifields, we quantified the orientation between pre- and post-treatment coding axes using the cosine similarity of their weight vectors. Cosine similarity ranges from −1 to 1, with values near 1 indicating similarly oriented coding axes, values near 0 indicating orthogonal axes, and values near −1 indicating oppositely oriented axes. Accordingly, pre/post cosine similarity was markedly reduced by ear plugging compared to sham treatment (mean: 0.08 versus 0.51. 95% CI: -0.1 - 0.21 versus for 0.44 – 0.59 for ear-plugged and sham mice, respectively) and was on average similar from classifiers trained on shuffled trial labels (Figure 7C). To determine if this low similarity simply reflected variability in estimating the classifier axis from finite trials, we generated independent coding axes within each pre or post-treatment session by training classifiers on separate split halves of trials and measuring their cosine similarity. These within-session axes remained substantially aligned in both ear-plugged and sham cohorts (Figure 7C. Mean: 0.44 versus 0.55. CI: 0.37 – 0.50 versus 0.47 – 0.63 for ear-plugged and sham mice, respectively). A two way, mixed design ANOVA revealed main effects of treatment group (F(1,20) = 21.65, p<0.001), cosine comparison (F(2,40) = 86.26, p<0.001) and a treatment group x cosine comparison interaction (F(2,40) = 18.82, p<0.001). Thus, the near-zero alignment between pre- and post-plug classifiers cannot be explained by instability of the decoder itself. Instead, the data further argue that monaural hearing loss reorients the population activity carrying hemifield information to a nearly orthogonal axis, while nevertheless allowing spatial information to remain linearly decodable.

### 2.8 Distinct neurons contribute to spatial coding accuracy in pre- and post-plug conditions

Shell IC population activity may retain a surprising degree of sound-location information after monaural hearing loss, albeit via activity patterns distinct from conditions where the full complement of binaural cues is available. To what extent might downstream readouts rely on distinct neuron groups for binaural vs monaural decoding of sound location? We addressed this question by training multinomial logistic-regression classifiers to decode absolute sound-source location from population activity in the pre- and post-treatment conditions (Figure 8A). Similar to binary classifiers, multinomial classifiers trained on pre-plug or post-plug data performed significantly above chance level when tested on held out trials from within the same condition, but they did not generalize across conditions; classifiers trained on sham-treated data readily generalized across conditions (Figure 8B). A two-way ANOVA revealed significant main effects of treatment condition (plug vs. sham, F(1,20)=4.54, p=0.046), classifier training condition (F(3,60)=45.42, p<0.001), and a significant treatment x training condition interaction (F(3,60)=5.16, p=0.034). Moreover, the mean absolute error (MAE) was markedly higher in cross-compared to within condition testing (Fig 8C), such that classifiers tested across conditions predominantly classified sounds as originating from the contralateral, open ear (Figure 8D) ^18^. A two-way ANOVA on the MAE data revealed significant main effects of treatment condition (plug vs. sham, F(1,20)=9.17, p=0.007), classifier training condition (F(3,60)=56.88, p<0.001), and a treatment x training condition interaction (F(3,60)=14.98, p<0.001).

**Figure 8:**
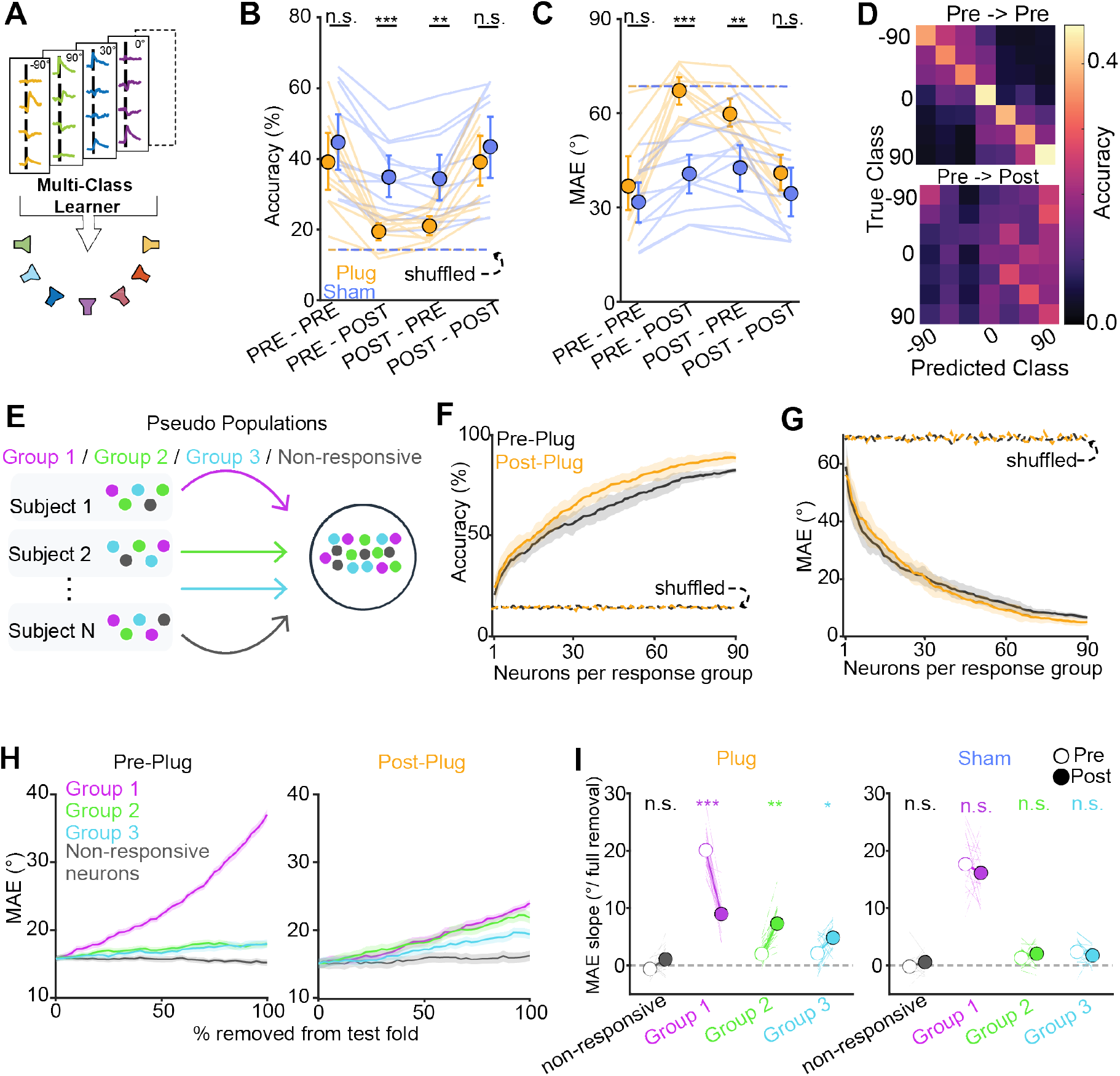
Neuron Groups 1-3 differentially contribute to spatial readout before and after ear-plugging. A) Cartoon. Multinomial classifiers are trained to classify sound location from single trials. B) Classifier accuracy is plotted across conditions for ear-plugged (yellow) and sham-treated (blue) mice. Bubbles and error bars are mean ± 95% CI from 1,000 bootstraps across subjects. Lines are individual subjects. ***, p<0.001, **, p<0.01. n.s., not significant. C) MAE across conditions for ear-plugged and sham-treated groups. Color scheme is same as B. Of note is the large increase in error magnitude when classifiers are trained and tested across conditions. D) Confusion matrices for classifiers trained and tested on pre-Plug datasets (top) and trained on pre-plug, tested on post-plug datasets (lower panel). E) Cartoon diagram of pseudopopulation approach. F) Accuracy as a function of neurons for pre- and post-plug classifiers (black and yellow, respectively). G) same as (F) but for MAE. H) MAE as a function of fraction of neurons removed from the testing fold for classifiers trained on pre- and post-plug pseudopopulations (left and right, respectively). I) Left: The slope of linear fits to the pseudopopulation data in panel H. Open and closed symbols are the mean of pre- and post-plug conditions, respectively. Lines are individual pseudopopulations (n =20). ***, p<0.001. **, p<0.01. *, p<0.05. n.s., not significant. Right: Same as panel, for pseudopopulations built from sham experiments.

To distinguish a global loss of spatial signals from a redistribution of information across neuronal response groups, we constructed pseudo-populations containing equal Group 1, Group 2, and Group 3 neurons, and neurons that were not significantly sound responsive. Increasing population size improved accuracy and reduced MAE before and after plugging (Figure 8F,G), indicating that ear plugging did not substantially reduce the total spatial information available to a linear readout. To test if this preserved information was carried by different neuronal groups under binaural and monaural conditions, we trained classifiers on 20 separate pseudo-populations each containing k = 40 neurons each drawn from Groups 1, 2, 3, or non-responsive neuron (k = 160 neurons per pseudopopulation). We then progressively silenced each neuron group in the held-out trials at test time by setting the feature weights to 0. This manipulation measured the decoder’s dependence on each group’s activity without allowing compensation through retraining. Under pre-plug conditions, progressive removal of Group 1 neurons sharply increased the MAE at testing time, whereas removal of Group 2, Group 3, and non-responsive neurons were of less consequence (Fig 8H, Left). These results corroborate our findings that under pre-plug conditions, Group 1 neurons are the predominant drivers of neural population trajectory separation. However, MAE in post-plug conditions was comparably less sensitive to removal of Group 1 neurons, whereas removal of Group 2 and 3 neurons caused a greater degree of progressive increase in MAE compared to the pre-plug condition (Figure 8H, right). (Figure 8H, grey traces). To quantify group-specific removal sensitivity, we fit the relationship between the fraction of neurons removed and MAE; larger positive slopes indicate greater decoder dependence on that group. Following ear plugging, removal sensitivity decreased for Group 1 neurons (Figure 8I, left panel. Mean vs ± SEM: 20.11 ± 0.74 vs 8.99 ± 0.61 degrees/neuron removed, p<0.001, paired t-test). By contrast, removal sensitivity increased for Group 2 (2.03 ± 0.37 vs 7.32 ± 0.65, p <0.001) and Group 3 (2.16 ± 0.59 vs 4.87 ± 0.52, p = 0.01). Thus, compared to pre-plug, post-plug classifiers’ accuracy was more dependent on neurons from Groups 2 and 3. Importantly, this result was not simply due to general increase in sensitivity of post-plug classifiers to neuron ablation: removal sensitivity of non-responsive neurons was not significantly different across pre-or post-plug conditions (Figure 8H,I: -0.58 ± 0.29 vs 1.1 ± 0.48, p = 0.07), and removal sensitivities of all Groups were not significantly different across pre- and post-conditions when pseudo-populations were generated from sham-treated mice (Figure 8I, Right panel, all p > 0.05). Altogether, our results demonstrate a population-level mechanism that enables shell IC populations to rapidly adapt to abrupt loss of binaural cues, thereby providing a putative substrate to flexibly switch between spatial hearing contexts.

## 3 Discussion

We have shown that in the mammalian shell IC, spatial auditory representations persist despite a dramatic and sudden reduction of binaural input. These results are surprising given that ipsilateral ear-plugging typically broadens single neuron spatial receptive fields in various brain regions ^30;101;102^ (but see ^31;33;34^). Our data reconcile these prior findings by showing that abrupt monaural hearing loss only degrades spatial tuning in a minority population of ON responsive Group 1 neurons. By contrast, the more numerous sound inhibited, OFF responsive Group 2 neurons maintain spatial selectivity following monaural hearing loss and become dominantly informative of sound source location after binaural cues are altered. These data provide a mechanistic explanation for how a sensory system maintains non-overlapping and context-specific spatial representations that can be flexibly recalled depending on peripheral cue availability.

Spatial tuning of Group 2 OFF responses may arise via multiple non-binaural mechanisms such selectivity to monaural spectral cues ^30;69;70^, or possibly, non-monotonic “O-shaped” level dependence of acoustic inputs ^103^. Alternatively, we cannot rule out that ear plugging causes a rapid (< 2 hours), functional remapping of binaural inputs that is independent of extensive experience. Indeed, even structural plasticity can within hours of altering peripheral inputs in adult mammals ^104–106^. However, this explanation assumes that any such remodeling is target-specific and first occurs in Group 2 neurons while minimally impacting intermingled neurons of distinct response classes. No such mechanism has ever been described to our knowledge, and is thus a less parsimonious explanation than assuming that spatial selectivity in Group 2 neurons persists in absence of typical binaural input.

### 3.1 Rotational dynamics enable co-existing and context-specific spatial representations

At the population level, binaural and monaural representations occupied largely non-overlapping coding subspaces with distinct geometries. Yet, despite this profound reorganization, neural responses to different sound-source locations remained separable within each subspace, supporting comparable spatial decoding across hearing conditions. Thus, monaural hearing loss did not simply degrade an existing spatial representation, but instead reoriented the population dimensions along which spatial information was encoded. Similar rotational dynamics are well established in motor systems, where low-dimensional neural activity cycles through orthogonal but functionally related subspaces during movement preparation and execution ^94;99;107^, or when similar movements are executed over time ^93^ and across different behavioral tasks ^108^. Such re-organization allows the same neural population to support distinct low-dimensional computations while minimizing interference, efficiently separates of past and present information ^109^, and may potentially operate on sub-second timescales ^110;111^. More broadly, these results highlight how context-dependent changes in the geometry of activity can implement distinct computations within a shared neural circuit. Interestingly, early experiments in the barn owl external IC found that adaptive remapping of auditory space during prism-rearing did not erase the original auditory spatial map, but instead rendered it functionally latent through GABAergic inhibition ^112^. Given the suggested homology between the mammalian shell IC and avian external IC ^113;114^, these findings imply a common principle of context-dependent spatial coding: The availability of peripheral cues determines which of several coexisting representations is functionally expressed at a particular moment in time.

### 3.2 Implications for spatial cue plasticity following monaural hearing loss

Acute monaural hearing loss profoundly disrupts sound localization, yet humans and animals can recover spatial hearing given appropriate training and experience ^48;49;115^; this perceptual learning has traditionally been attributed to the recalibration of altered binaural cues or increased reliance on monaural spectral cues ^15;19;116;117^. Interestingly, listeners who learn to localize sounds under both binaural and altered conditions can flexibly switch between strategies with minimal re-learning or interference ^21;38;118;119^. Our results suggest a potential neural substrate for this flexibility. Within hours of monaural hearing loss, the auditory midbrain maintains a spatially informative code, largely orthogonal to the original binaural representation and dominated by OFF rather than ON responsive neurons. These results conspicuously dovetail with human sound localization studies reporting prolonged reaction times with unilateral hearing loss ^120;121^, as well as delayed cortical responses to acoustic spatial changes even in listeners with near-normal localization accuracy ^122^. We therefore propose that recovery from monaural hearing loss may rely less on repairing a degraded binaural representation than on down-stream circuits learning to employ a newly informative, OFF-dominated population code. In this framework, the neural representation necessary for accurate spatial hearing emerges rapidly, while the rate-limiting step to perceptual recovery reflects the slower process of learning to exploit it.

## 4 Methods

### 4.1 Animal subjects and handling

All procedures were approved by the University of Michigan’s Institutional Animal Care and Use Committee and carried out in accordance with the NIH’s guide for the care and use of laboratory animals. We used a total of 16 normal hearing mice 10-14 weeks at the time of surgery), F1 offspring of C57.Bl6/J x CBA/CaJ breeders bred in our colony. N = 15 mice were employed for imaging experiments and one mouse was used solely for ABR testing ear plug efficacy. Mice were single-housed with visual and olfactory contact to neighboring animals at a reversed 12/12-hour dark-light cycle, food and water were provided *ad libitum*. Cages were equipped with standardized enrichment (running wheels, shelter, and two different forms of nest-building material) to reduce stereotypic behavior and stress. Experiments were conducted during the dark period. Handling by the experimenter began at least 10 days following surgery and comprised 3-10 sessions of habituation to the Plexiglas seating tube in which mice sit during head fixation. During these sessions, mice were allowed to explore the tube in the cage, followed by sessions of exploring the tube in the hand of the experimenter. In the last 2 sessions, mice were carefully fixated by holding the head bar for a few seconds.

### 4.2 Surgeries

Surgeries were conducted under isoflurane anesthesia between 10-14 weeks of age as described previously ^50;51^. The viral construct for the expression of the Ca^2+^ indicator GCaMP6f (n = 2, pAAV1.Syn.GCaMP6f.WPRE.SV40, Addgene, titer order of magnitude 10^12^) or GCaMP8s (n = 13, pAAV1.Syn.GCaMP8s.WPRE, Addgene, titer order of magnitude 10^12^) was pressure ejected at 4 sites ∼200 µm below the dura (25 nl each; 100 nl total) across the medial lateral axis of the IC using an automated injection system (Nanoject III, Drummond). A custom-made titanium head bar was placed on the skull and encased in dental resin, keeping the pinnae fully exposed for naturalistic binaural and monaural spectral cues. Buprenorphine (0.03 mg/kg, PAR pharmaceutical) was administered subcutaneously at the end of the surgery to reduce pain, and mice were allowed to recover in a clean cage on a heating pad for approximately 1 hour. Mice were provided with a purified high energy dietary supplement (DietGel Boost, Clear H2O) to support recovery. Following 24 and 48 hours, mice received additional injections of carprofen. Well-being and surgical sites were closely monitored for 7 days following surgery.

### 4.3 Sound delivery system

We developed a moveable speaker system to deliver spatial sound stimuli. Experiments using 7 distinct speaker positions (Figures 1-3) employed a servo motor (2000 Series Dual Mode Servo 25-2, Torque or Teknic CPM-MCPV-2310S) equipped with a 30 cm arm (33-hole aluminum flat beam, Actobotics), carrying the speaker (XT25SC90-04, Peerless by Tymphany) at the distal end via a custom 3D printed mount. The servo was centered under the head of the head-fixed animal, enabling 180° speaker rotation within the frontal horizontal field. Servo movement was controlled via pulse-width modulation signals generated in MATLAB and delivered via an analog headphone channel on a high-fidelity sound card (Fireface UFX+, RME). The commands were fed into a custom-made signal conditioning circuit to smooth and amplify the output signal to 5V. Duty cycles for each position were carefully calibrated to an accuracy of 1-2°. For the 2000 Series servo, trial-to-trial positional accuracy was additionally controlled using a “hardwired” closed loop control of speaker position using a microcontroller (Arduino UNO, Arduino AG) and a photo interrupter mounted on a 3D printed slotted disc, with each slot corresponding to the specific speaker angles used in a particular imaging session. Trial-to-trial positional accuracy of the Teknic motor was precise enough to obviate the need for closed loop stabilization. Servo movements occurred selectively during the inter-trial interval, at least 15 s prior to data acquisition of each trial. Position commands were generated in MATLAB and delivered via a Bpod analog output module (Sanworks) without additional signal conditioning.

Sound stimuli (4-35 kHz frozen broad-band noise with 5 ms on/offset cosine ramps, 500 ms duration) were generated in Matlab (192 kHz sampling rate) and presented at 65 dB SPL via a high-fidelity sound card (Fireface UFX+, RME) and 200 W power amplifier (SLA-2, ART). The speaker was positioned on each trial between -90° to 90° (zeroed on midline), with most experiments employing 7 positions [-90° -60° -30° 0° 30° 60° 90°]. A subset of experiments tested additional positions at ± 45° and ± 15°; these positions were omitted from analyses of ear-plug effects in Figures 3-8. The speaker was calibrated at each position using a 1/4” pressure-field microphone (CCLD pressure-field, Type 4944-A, Bruel & Kjaer). The microphone was positioned using a custom 3D-printed attachment for the head bar holders which centered the microphone at the approximated center of the animal’s head, with the microphone membrane perpendicular to the horizon.

### 4.4 2-photon Ca^2+^ imaging

Experiments were performed in darkness in a table-top, sound-attenuating chamber. Mice were placed in a Plexiglas tube and head fixed on a custom-built rig. The experimental setup (microscope, head-fixation system, and sound delivery system) was placed on an optical table and surrounded by a sound attenuating double-walled booth, lined with acoustic foam. Data were collected with a Janelia-style MIMMs or Sutter MOM two-photon microscope system. Movies were collected at a frame rate of 30 Hz and a resolution of 512×512 pixels using a resonance scanner, a 16x water immersion objective (Nikon, 0.8 NA, 3 mm working distance), and a GaAsP photomultiplier tube (Hamamatsu Photonics). GCaMP was excited at 920 nm using a Titanium-Sapphire laser (Chameleon Ultra 2, Coherent) which was positioned on the air table outside of the booth. Each imaging trial consisted of 1-3 sound presentations at a specific angle, with a minimum of 5 s between each sound presentation. The speaker was moved to a new position during a ∼15-20 s inter-trial interval. 5-10 s elapsed following motor movement and sound delivery on the subsequent trial to minimize temporal overlap between sound responses driven and prior movement.

### 4.5 Ear plugging

Mice underwent two separate imaging sessions (pre- and post-plug) on the day of ear-plugging. To reduce stress, we waited at least 1.5 hours between pre-plug session and ear plugging. Mice were anesthetized using isoflurane and placed on the bite bar and heating pad of a rotatable stereotaxic frame without the support of ear bars. Here, anesthesia was maintained between 1-1.5 % isoflurane due to the non-invasive nature of the procedure and to achieve a rapid recovery. Lubricant was applied to the mouse’s eyes, and bite bar and gas mask were carefully rotated while constantly adjusting the posture of the animals’ body until the left ear (ipsilateral to the IC window) was exposed and faced upwards. Mice received a shot of carprofen (2.5 mg/kg s.c.) to reduce any potential discomfort associated with plug placement and thus to reduce scratching and grooming behavior to avoid early plug loss. Both pinna and distal portion of the ear canal were cleaned with ethanol and fur that reached into the ear canal and pinna was removed. The appearance of the ear canal and the tympanic membrane were evaluated using a small digital otoscope (Teslong). A standard human earplug (Mack’s Ultra Soft Foam Earplugs – NRR 33 dB, McKoen) was cut in a cone-like shape with an outside facing diameter of ∼5 mm and a maximum length of 3 mm. The plug was superficially wiped with ethanol, dried, compressed using forceps, carefully inserted into the ear canal, and released. Following expansion, the plug was checked for a tight seal around the entire edge before a skin-friendly 2-component silicone rubber (BodyDouble Fast, Smooth-On Inc.) was applied to cover the foam plug. The mouse was rotated back and then remained on the frame to allow the hardening of the silicone rubber (∼7 min). Post-plug imaging data were collected following a minimum of 1.5 hours recovery. To ensure that any change in spatial responses were due to monaural hearing loss rather than anesthetic delivery or representational drift ^123–125^, a separate control group of mice underwent a sham treatment, with similar anesthetic induction, stereotaxic placement, and imaging regime as described above but were not fitted with an earplug.

We tested the efficacy of our ear plug approach by measuring auditory brainstem responses (ABR) in two of the mice of the experimental group and one naïve mouse without a head bar. ABRs were obtained via an EPL Cochlear Function Test Suite (Eaton-Peabody Laboratories, Massachusetts Eye and Ear, as described in ^126^). Mice were anesthetized with ketamine (initial 10 mg/ kg, maintenance 2.5 mg/kg) and xylazine (initial 0.083 mg/kg, maintenance 0.01 mg/kg). Anesthesia depth was confirmed via toe pinch and anesthetic maintenance doses were provided as needed. Subcutaneous needle recording electrodes were placed behind the ear, and a reference electrode was placed at the vertex. A ground electrode was additionally placed at the tail. Electrode signals were bandpass filtered (300 Hz to 3 kHz) and amplified 10,000-fold. Sound was monaurally delivered directly into the ear canal (unplugged ears) or near field in front of the plug. Recordings were performed using National Instruments input/output boards hardware. To determine hearing thresholds at different sound frequencies, tone pips (5 ms, 0.5 ms cosine ramps at onset and offset) were presented at 8, 16, and 32 kHz. The intensity was varied between 10 and 80 dB SPL in 5 dB steps. An automatic online artifact rejection algorithm discarded trials with muscle potential artifacts according to a preset threshold. ABR data were recorded continuously and saved for offline analyses. For every stimulus at least 400 artifact free trials were recorded. Thresholds at individual sound frequencies were determined by eye from the average stimulus-aligned traces at each frequency as the lowest level that evoked an ABR. If no response could be evoked up to 80 dB SPL, threshold was set to 80 dB.

### 4.6 Imaging data analysis

We used the Python version of Suite2p ^127^ to motion correct the raw data and extract fluorescence time series from regions of interests (ROIs) corresponding to individual IC neurons. Datasets were manually curated to exclude overlapping ROIs. Raw fluorescence traces were converted to ΔF/F by dividing the fluorescence by the mean baseline intensity for each trial (1 s prior to sound onset; 30 frames), and peak amplitudes were measured as the average of ± 1 frames around the maximum or minimum ΔF/F value during sound ON or sound OFF periods. The surrounding neuropil signal was scaled by a factor of 0.7 and subtracted and traces were smoothed using a 5-frame gaussian kernel. ΔF/F traces were then further analyzed using custom MATLAB code (version 2021a, 2022b, and 2023a, and 2024b). To determine significantly responding ROIs, we used a bootstrapping procedure based on the trial-by-trial autocorrelation of ΔF/F waveforms on similar trial types ^51–53^. Briefly, the average correlation of each matching pair of trials with the same stimulus is compared to a randomly sampled signal from the same trials 10000 times. The p-value is computed as the fraction of these randomly sampled signals with greater signal autocorrelation than the real data. P-values were then corrected for multiple comparisons using the Bonferroni-Holm method.

To track the activity of neurons across ear-plugging and sham treatments, ROIs of both pre and post conditions were registered using a probabilistic, automated approach that quantitatively evaluates registration accuracy (CellReg) ^87^. Registration accuracy was visually inspected and curated for each session.

### 4.7 Clustering Analysis and Neuron Group Identification

To identify unique neuron groups that could account for the diversity of shell IC temporal response profiles, we first averaged each neuron’s fluorescence time series across all tested spatial positions, resulting in a nFrames x nNeurons matrix. We then reduced the dimensionality of the matrix using PCA. The first 20 principal components were subsequently retained for k-means clustering, using k = 3 clusters as indicate by silhouette score and WCSS analyses. Heatmaps in Figure 1 were generated by dividing trials into odd and even subsets, sorting neurons by the peak response time measured from odd-trial averages, and displaying their mean responses across even trials.

### 4.8 Orientation Selectivity Index

To measure the degree of spatial tuning in each neuron, We calculated an orientation selectivity index (OSI) similar to previous studies ^128^

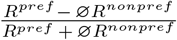

where R^pref^ is the maximum or minimum response (peak ΔF/F) at the most preferred angle and ∅R^nonpref^ is the mean response at all other positions. For OSI measurements in Figure 2, we included all significantly responsive neurons in Groups 1-3. For comparison of changes in OSI values in Figure 3, we only considered neurons that could be tracked across sessions, were significantly sound responsive in the pre-plug (or pre-sham) condition, and remained significantly responsive post-manipulation. 104/514 and 103/733 of neurons in ear-plugged and sham groups respectively became unresponsive post-manipulation and were excluded. This “ROI dropout” did not differ significantly between ear-plugged and sham treatment, or across neuron groups. A mixed effects model revealed no significant main effect of treatment condition (F(1,59)=1.88, p=0.18), no significant main effect of neuron group (F(2,59)=0.65, p =0.52), and no treatment condition x neuron group interaction (F(2,59)=0.56, p=0.57). Thus, the occasional loss of significant sound responses in our conditions is unlikely due to ear-plugging.

### 4.9 Neural trajectory analyses

To quantify how time-varying population activity is altered by monaural hearing loss, we constructed neural trajectories from matched pre- and post-plug or sham sessions using all neurons that could be reliably tracked across conditions, including non–sound-responsive neurons. Trial counts at each sound location were equalized by sub-sampling to the smallest number of trials available per mouse. For each neuron, ΔF/F responses were concatenated across pre- and post-sessions, z-scored using the mean and SD estimated jointly across pre- and post-manipulation trials, and projected onto a common low-dimensional basis obtained via PCA. We retained the first 15 PCs for analyses in Figures 4C-D. This captured 58.3 ± 2.7% and 59.0 ± 2.6% of the variance in plug and sham mice, respectively, thereby limiting contributions from low-variance dimensions. We confirmed that 15 PCs remained below the effective dimensionality of the datasets by calculating the Participation Ratio for each experiment as 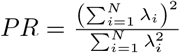 which suggested a mean effective rank of ∼20 for these datasets (Supplemental Figure 6C).

To estimate differences between pre- and post-plug population dynamics, we computed cross-validated Mahalanobis (crossnobis) distances ^129;130^. For each experiment, trials from both pre- and post-plug sessions were randomly split into two equal halves (A and B). Within each split, we averaged the time series separately for each sound location, and projected the resulting activity onto the PCA basis. Independent pre-to-post difference vectors were then calculated for the two halves, D_A_ and D_B_, defined as

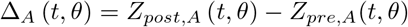

and

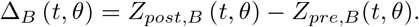

The cross validated dissimilarity then was calculated at each time point as 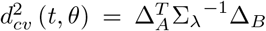. The noise covariance matrix Σ_*λ*_ was estimated in PCA space from residuals obtained after subtracting the corresponding session- and sound-location-specific mean response. Residuals were pooled across pre/post-plug sessions, sound locations, trials, and time points, and the covariance estimate was regularized by fixed linear shrinkage toward a scaled identity matrix (l=0.01). The procedure was repeated 30 times with A/B trial assignments randomly redrawn, and distance estimates were averaged across these 30 repetitions. Because crossnobis is an unbiased signed dissimilarity estimate, individual values could be negative. Scalar trajectory-difference measures were then obtained by summing the resulting time courses over the combined sound-ON and sound-OFF interval. Null distributions were generated from 1,000 permutations in which sound-location labels were shuffled independently within the pre- and post-manipulation sessions before projecting the data onto a fixed basis and calculating the distance between sound location trajectories.

### 4.10 Procrustes Transformation

For each mouse, neurons matched across pre- and post-treatment sessions were retained, and trials were averaged separately for each sound location and treatment condition. The resulting population responses were concatenated across time, sound location, and treatment conditions to form a (2*SoundLocationTime*) × *Neurons* matrix. A single PCA basis was fit to the pooled pre- and post-treatment condition means using MATLAB’s pca() function, and neural trajectories for each sound location and treatment condition were constructed by projecting trial-averaged and mean-centered population activity onto the first two PCs.

We quantified changes in trajectories using MAT-LAB’s procrustes() function, which finds the transformation that best superimposes post-plug trajectory *Y* onto pre-plug trajectory *X*. The transformation is defined according to *X* ≈ *bY T* + *c*, where *T* is a rotation matrix, *b* is an isotropic scaling factor, and *c* is a translation vector. To this end, trials were randomly divided into two equal halves (A and B) and each half was averaged separately per sound location and treatment condition. For each of 100 repetitions, across-session transformation variables (absolute rotation angle *T*, scale factor *b*, and translation magnitude *c*) were calculated as the mean of the PRE-A versus POST-A and PRE-B versus POST-B alignments for each sound location trajectory. Within-session controls were calculated as the mean of the PRE-A versus PRE-B and POST-A versus POST-B alignments, providing an estimate of transformations expected from sampling variability alone.

### 4.11 Representational Dissimilarity Analyses

For analyses in Figures 5, the representational separation between all sound-locations was measured in the full neural space without dimensionality reduction. In each pre and post-treatment session, single trials at each of the seven sound locations were randomly divided into two non-overlapping subsets, averaged, and distance timecourses were calculated for each of the 21 unique sound-location pairs using the crossnobis measurement described above (30 independent redraws). The distance time course was divided by the number of matched neurons in each recording to account for differences in population size, and averaged across all sound location pairs. Peak dissimilarity was calculated as the mean of the maximum dissimilarity frame and its two immediately adjacent frames.

For analyses in Figures 5E–H, single trials at each of the seven sound locations were randomly divided into four non-overlapping subsets. Two subsets were used to estimate the upper RDM triangle, with the other two subsets were used to estimate the lower triangle independently. For each RDM entry, the crossnobis measurement between sound locations in full neural space was calculated as above, and a scalar value was generated by summing across imaging frames beginning at sound onset until 1 s after sound termination. Trial partitioning was repeated 30 times and the resulting RDMs were averaged across repetitions, producing a 7 × 7 RDM containing independent estimates of the dissimilarity between every pair of sound locations. Diagonal entries representing cross-validated self-distances were excluded from subsequent summary and correlation measurements.

Summary statistics were quantified by calculating the averaging entries from both RDM triangles. Percentage change from PRE was calculated as

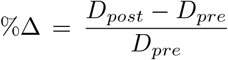

Where *D*_*pre*_ and *D*_*post*_ denote the mouse’s mean off-diagonal dissimilarity before and after treatment, respectively. Correlations between pre and post-treatment RDMs were calculated independently for the off-diagonal entries of the upper and lower triangles using Spearman’s rank, and then averaged to obtain one correlation value per mouse. Values from plug and sham mice were compared using separate two-sided Welch’s t-tests.

### 4.12 Pseudo-population Analyses

Pseudo-populations in Figures 5I-M were constructed using neurons matched between pre- and post-treatment sessions. One Group 2 neuron with an unusually large post-plug response relative to the pre-plug condition was excluded, to prevent its repeated sampling from disproportionately influencing the pseudo-population analysis; Group 2 effects were larger in magnitude but similar in direction when this neuron was included. For every pseudo-population draw, k = 10 neurons were sampled from a single response group, with eligible neurons drawn as evenly as possible across mice. To this end, one neuron was drawn from each mouse until the required 10 neurons had been selected. If the number of neurons could not be divided equally among mice, the randomized mouse order determined which subject supplied the remaining neurons. Mice lacking eligible neurons in a particular response group were skipped for that group; if a mouse’s available neurons were exhausted, the remaining neurons were supplied by other mice on that draw. Mouse order and neuron selection were independently randomized for every draw. Sampling was without replacement within a draw and performed separately for each neuron group, though neurons could be selected again in subsequent draws. Pseudo-trials were assembled independently for each neuron, treatment condition, and sound location. The number of pseudo-trials in each condition and sound location was determined from the median number of available trials across the neurons selected for that draw. For every neuron, the required trials were randomly selected without replacement when sufficient trials were available, and with replacement only when a neuron had fewer trials than the requested pseudo-trial count. The chronological index of trials within a recording session were shuffled prior to selection, thus limiting noise correlations across pooled neurons due to trial order ^131;132^. The dissimilarity timecourses and RDMs were then calculated and summarized as described above, separately for each neuron group and pseudopopulation.

### 4.13. Subspace alignment

We quantified subspace alignment of population activity across hearing conditions as described in Elsayed et al., 2016. Analyses were restricted to neurons matched between pre- and post-manipulation sessions. For each experiment, Fluorescence traces were normalized as described above for neural trajectory analyses. For each pre/post condition pair (i.e., pre- and post-plug), traces were averaged across trials separately for each neuron and each sound location, and the across-location mean of that pre/post condition was subtracted at each time point. The resulting traces were row concatenated and combined into a matrix of size N x LT, corresponding to neuron × sound location-time in the particular pre/post condition. Data matrices for each condition were decomposed separately using singular value decomposition. The first six components defined the pre and post subspaces. We then measured the variance in each activity matrix captured by its native basis and by the basis obtained from the other session. Subspace overlap was quantified using the alignment index:

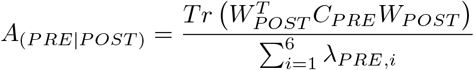

where *W*_*post*_ contains the first six post-manipulation PCs, *C*_*pre*_ is the pre-manipulation activity covariance matrix, and *λ*_*PRE,I*_ are its ordered eigenvalues. The reverse-direction index was calculated analogously, and the two directional indices were averaged for group summaries. Null distributions were generated separately for each projection direction using 10,000 random six-dimensional subspaces sampled according to the covariance structure of the combined pre- and post-manipulation activity, following Elsayed et al. (2016). One-sided empirical p-values were calculated as the fraction of random-subspace indices less than or equal to the observed index. An experiment was classified as significantly misaligned when either directional comparison reached p<0.025, accounting for the two projection directions.

### 4.14 Binary classifiers

For each mouse, only neurons matched across the two sessions were included as features for classifier training and testing. For each trial and neuron, activity was summarized as a scalar by integrating the ΔF/F fluorescence time series over a 2 s period following sound onset. Trials at all positions except the midline 0° trials were included in both training and testing. Before cross-validation, trials were balanced separately within the pre- and post-manipulation sessions by randomly downsampling each sound location to the smallest trial count available in that session. For each source training fold, each neuronal feature was z-scored using its mean and standard deviation across the source training trials. The resulting training-derived parameters were then applied unchanged to both the within-session and cross-session test trials ^133^. Linear support vector machine classifiers were trained to classify sound source hemifield using only non-midline sound location trials, via MATLAB’s sequential minimal optimization solver. The resulting classifier hyperplane is defined by an orthogonal weight vector w^T^ of size = neurons and an intercept term *b*. Via the linear equation f(x) = w^T^x + b, the projection f(x) classifies single trial features in the vector x as belonging to a binary category by the sign of f(x).

Classifier were evaluated across four train/test regimes: pre-to-pre, pre-to-post, post-to-pre, and post-to-post. Pre- and post-manipulation trials were independently assigned to five folds stratified by sound location. For each fold, a classifier trained on 80% of the pre-manipulation trials was tested on the held-out pre- and post-manipulation folds, and a classifier trained on four-fifths of the post-manipulation trials was tested on the held-out post- and pre-manipulation folds. Accuracy was calculated separately at each sound location and then averaged across locations so that each location contributed equally. The complete cross-validation procedure was repeated five times with newly generated fold assignments, and accuracies were averaged across repetitions. Shuffled-label controls used the same pipeline, except that the source training labels were shuffled independently within the pre- and post-manipulation training folds while the test labels were preserved. This procedure was repeated for 100 permutations.

Classifier alignment was quantified as the cosine similarity between the pre- and post-session SVM weight vectors, calculated as

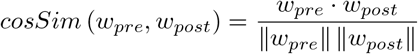

Values range from ™1 to +1, where +1 indicates parallel orientations, 0 indicates orthogonal orientations, and ™1 indicates opposing orientations. For each fold and repetition, we calculated the signed cosine similarity between the pre- and post-session classifier weight vectors after transforming the weights from their fold-specific standardized feature spaces back into raw integrated-ΔF/F coordinates. Cosine similarities were averaged across the five folds and five repetitions to obtain one mean alignment value per mouse. A null distribution was calculated from the weight vectors of the same 100 shuffled-label classifier sets used for the accuracy controls. Within-session reliability was assessed separately for the pre and post sessions using 100 repeated split-half resamples. For each resample, separate classifiers were fitted to two separate halves of the session’s trials, and cosine similarity was calculated between their weight vectors in raw integrated-ΔF/F coordinates. These split-half distributions provided benchmarks for within-session classifier alignment.

### 4.15 Multinomial classifier analyses

Single-trial activity from neurons matched between pre- and post-treatment sessions was summarized as described for the binary classifiers. Sound location was decoded using a multinomial logistic regression model, optimized with an L2 penalty on the neuronal feature weights; intercept terms were not penalized. For each trial, the classifier calculated a linear score for every sound location and converted the score into a class probability using the softmax function. The predicted location was the class with the greatest probability. Performance was quantified as classification accuracy and mean absolute error, defined as the absolute linear difference in degrees between the predicted and true sound locations. Correctly classified trials therefore contributed an error of zero. Classifier accuracy within and across treatment conditions, as well as shuffled label controls, were evaluated as described for the binary classifiers.

For pseudo-population analyses, we generated 20 pseudo-populations per treatment condition, containing 40 neurons from each of the three sound-responsive groups and the non-responsive neuron group. Neurons were sampled without replacement as described above (see Pseudo-population analyses), and we only included neurons which could be tracked across pre- and post-treatment conditions. To test how different neuronal response groups contributed to the classifier’s readout of sound location before and after plugging, classifiers were first trained on an intact pseudo-population’s pre- or post-treatment trials. Neurons belonging to a specific group were “removed” in the held-out test trials by setting the standardized test trial feature values to zero. Removed subsets progressed one neuron at a time, and were nested so that each level included all neurons removed at the preceding level. Removal sensitivity was summarized separately for every pseudo-population before and after plugging by calculating the slope of a linear fit between MAE and the fraction of neurons removed.

### 4.16 Statistics

Statistical comparisons were conducted in Matlab, Graphpad Prism, or Microsoft Excel using the procedures listed in the text. Significance level was set to alpha = 0.05 and all tests were two-tailed unless otherwise specified. Alpha was corrected for multiple comparisons when appropriate using the Holm-Bonferroni procedure. For ANOVA omnibus tests, p-values are reported after Geisser-Greenhouse correction unless otherwise noted. In Supplemental Figures 1 and 2, we tested if the distribution of preferred angles for Group 1 and Group 2 neurons differed from chance by generating 1,000 null distributions, uniformly sampled with replacement from the preferred angles present in that group. For each iteration, the observed distribution was compared to the null distribution using a Kruskal-Wallis test. The final reported p-value reflects the null-consistency score, i.e. the proportion of iterations in which the Kruskal–Wallis test was not significant.

## Author contributions

MMR, GLQ, and PFA designed the research. HLN, MMR, CMV, and ANF conducted experiments. GW collected auditory brainstem response data under the supervision of GC. MMR, GLQ, HLN, and PFA analyzed the data. MMR, GLQ, and PFA wrote the first draft of the paper. PFA wrote the final draft.

## Acknowledgements

We thank thank Stephen F. Wollgast, Dr. Rainer Beutelmann, and Deepak Dileepkumar for the support with sound delivery hard- and software; Dr. Michael T. Roberts for sharing sound delivery code and critical comments on the manuscript; Jordyn E. Czarny and Hannah M. Oberle for the support with mouse care. This work was supported by a Deutsche Forschungsgemeinschaft (DFG) Walter Benjamin Fellowship awarded to MMR (RO 6660/1-1:1) and a National Institutes of Health grants to R01DC019090 and RF1MH141903 to PFA.

## Declaration of generative AI and AI-assisted technologies in the manuscript preparation process

During the preparation of this work, the author(s) used OpenAI’s ChatGPT and Anthropic’s Claude to improve manuscript drafting, figure generation, and analysis. The author(s) reviewed and edited the output as needed and take full responsibility for the content of the published article.

## Supplementary Figures

**Figure S1:**
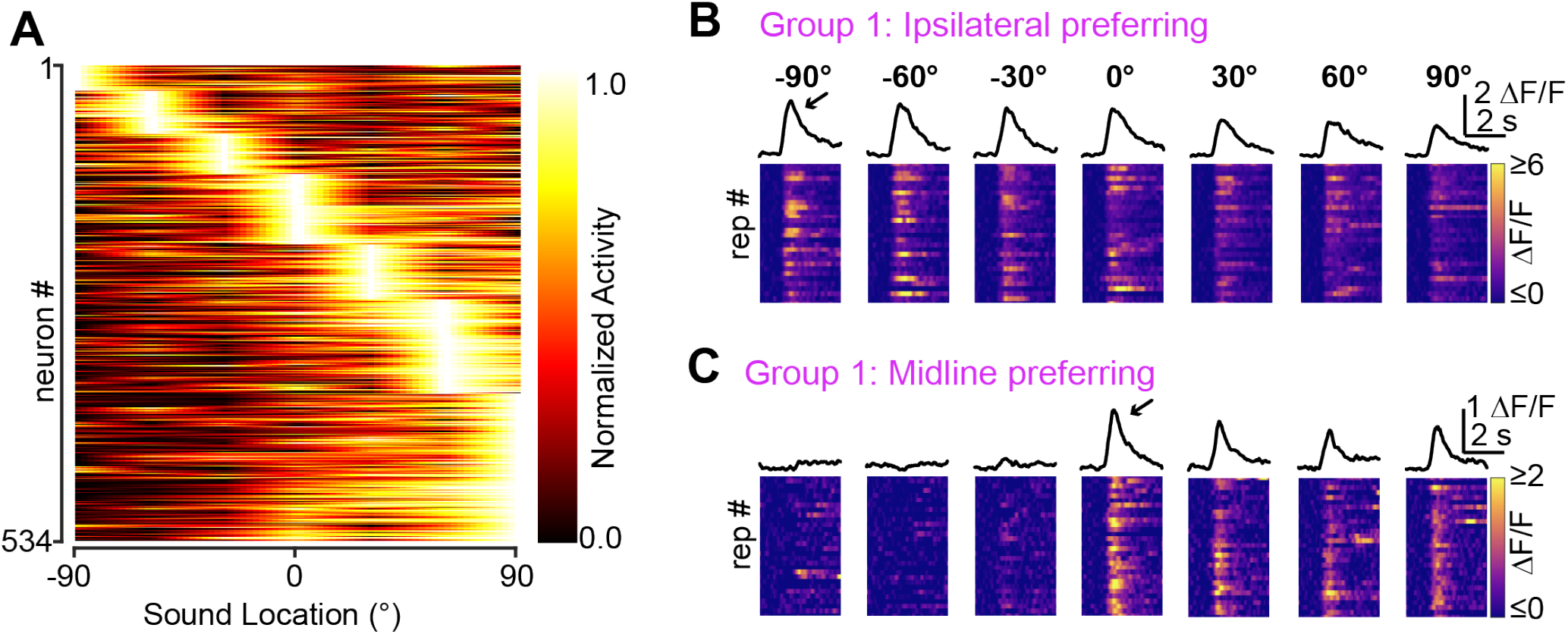
Spatial receptive fields of Group 1 neurons are predominantly, though not exclusively, contralateral. A) Summary heatmap of Group 1 neurons as a function of sound location. Neurons are row normalized and ordered by preferred location, revealing a contralateral bias in spatial tuning (p < 0.001, Kruskal–Wallis test compared to 1000 null distributions; see Methods). B) Example Group 1 neuron with peak response in the ipsilateral hemifield. C) Example Group 1 neuron with a peak response at the midline.

**Figure S2:**
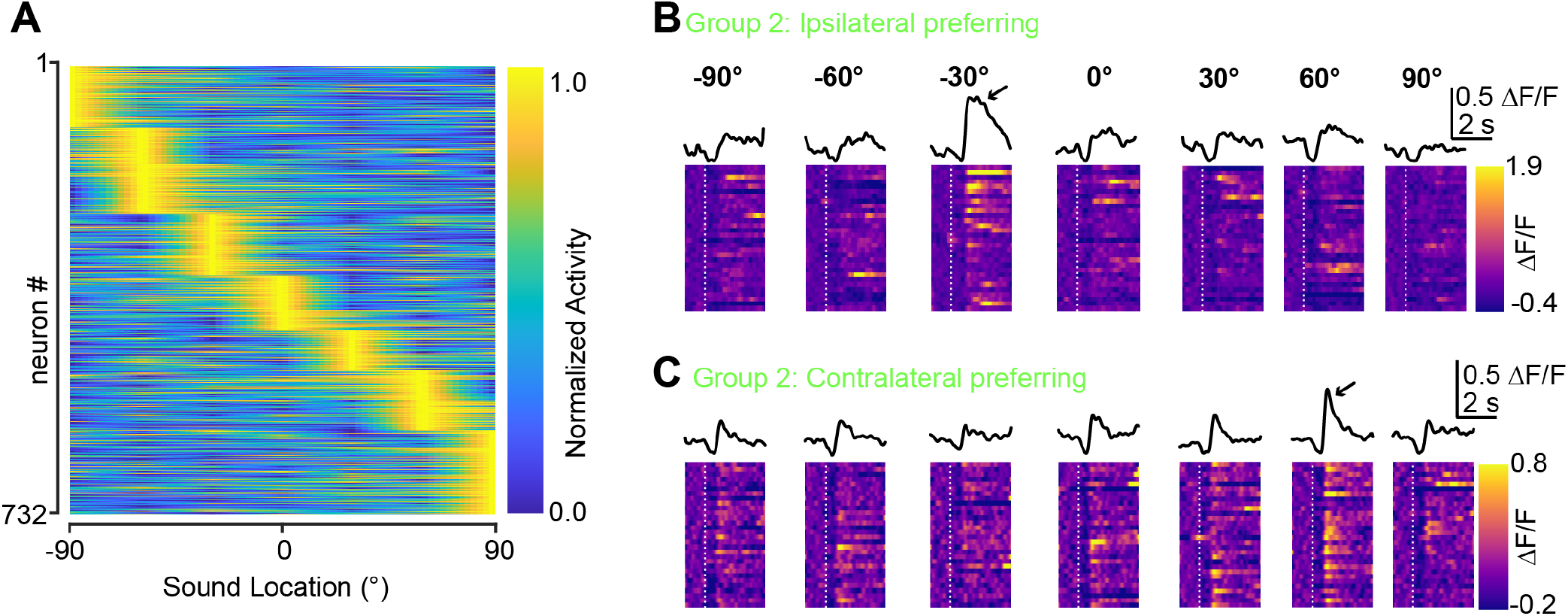
Receptive fields of Group 2 OFF responses tile the entire frontal azimuth. A) Summary heatmap of Group 2 neurons’ OFF responses as a function of sound location. Neurons are row normalized and ordered by preferred position, revealing roughly equal proportions of ipsi- and contra-lateral preferring neurons (p = 0.996, Kruskal–Wallis test compared to 1000 null distributions). B) Example Group 2 neuron with a preferred OFF response at -30° in the ipsilateral hemifield. C) Example Group 2 neuron with a preferred OFF response at 60° in the contralateral hemifield.

**Figure S3:**
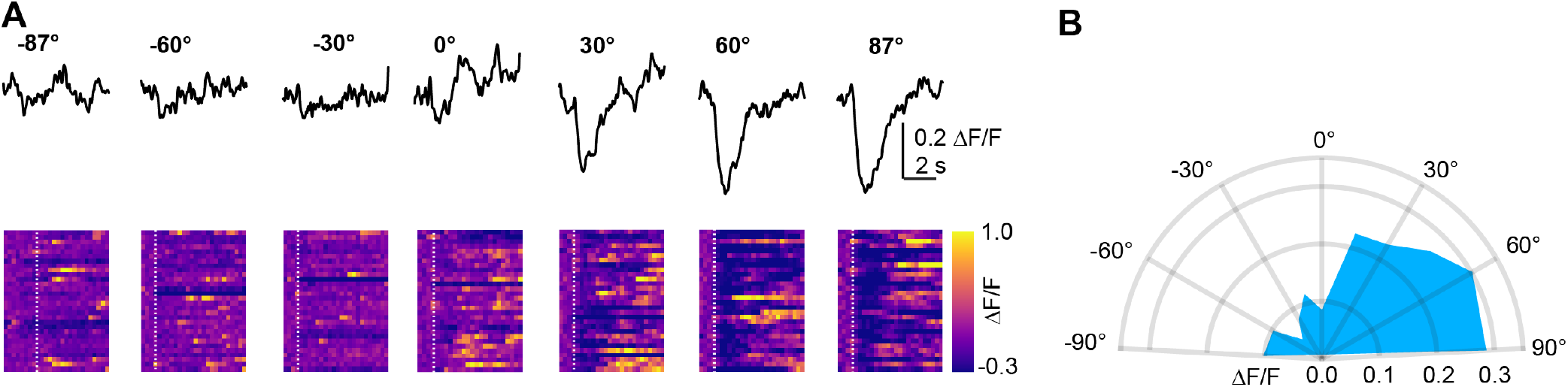
Example of directional inhibition in a Group 3 neuron. A) Group 3 neuron with sound inhibited responses evoked primarily by contralateral sounds. B) half-polar plot for the neuron in (A). The receptive field is constrained to contralateral space.

**Figure S4:**
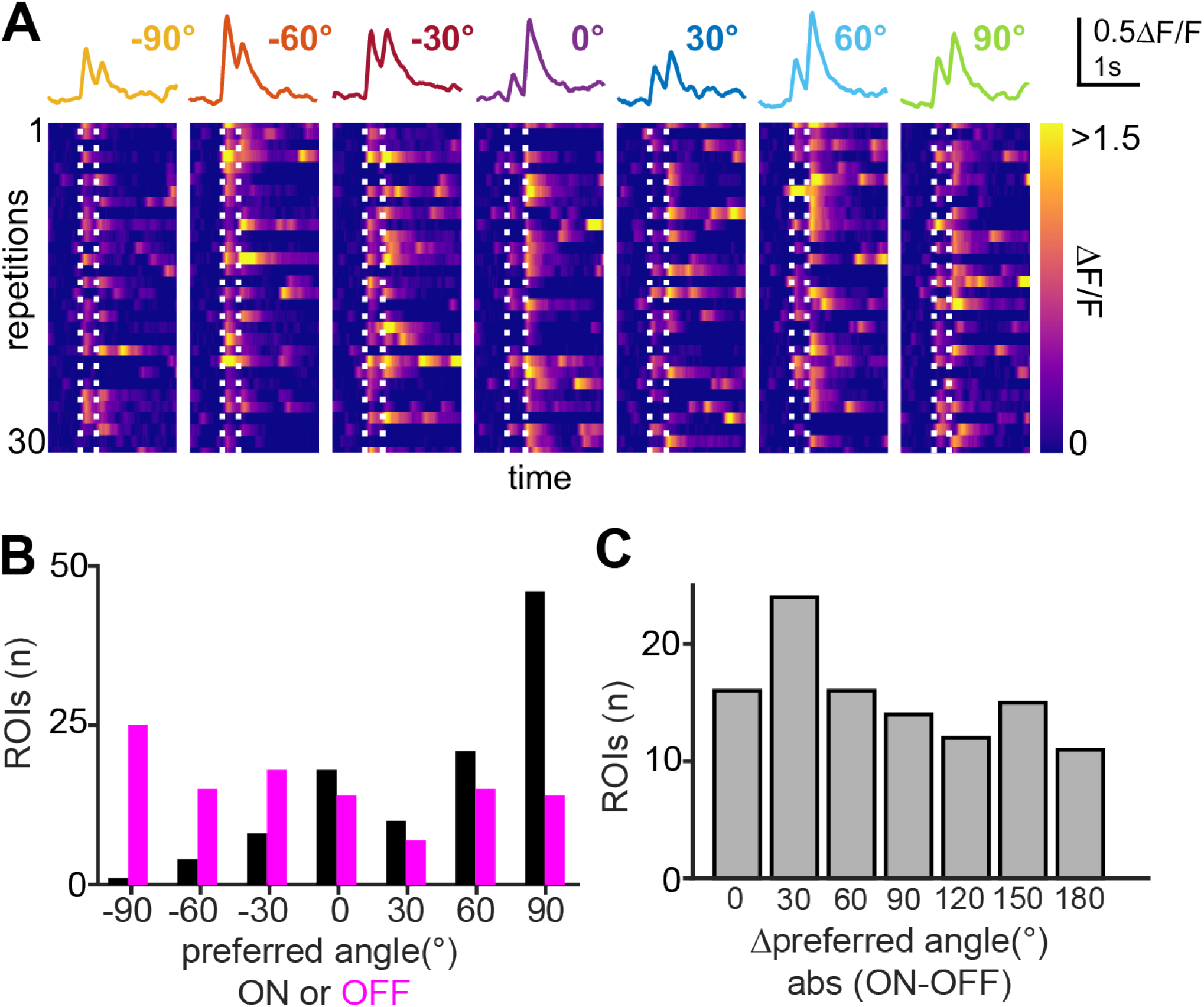
ON and OFF excitation in single neurons can have different receptive fields. A) Example neuron with ON-OFF excitatory responses. Top traces: Mean fluorescence response at each presentation angle. Lower panels: Heatmaps of individual trials. Dashed lines: sound onset and offset. B) Histogram of preferred ON and OFF excitatory responses for n = 71 neurons. The distributions were significantly different, two sample Kolmogorov-Smirnoff Test statistic: 0.42, p<0.001. C) Histogram of the absolute difference in preferred position of ON and OFF responses, measured within the same neuron. Of note is, most neurons have ON and OFF responses with distinct preferred locations.

**Figure S5:**
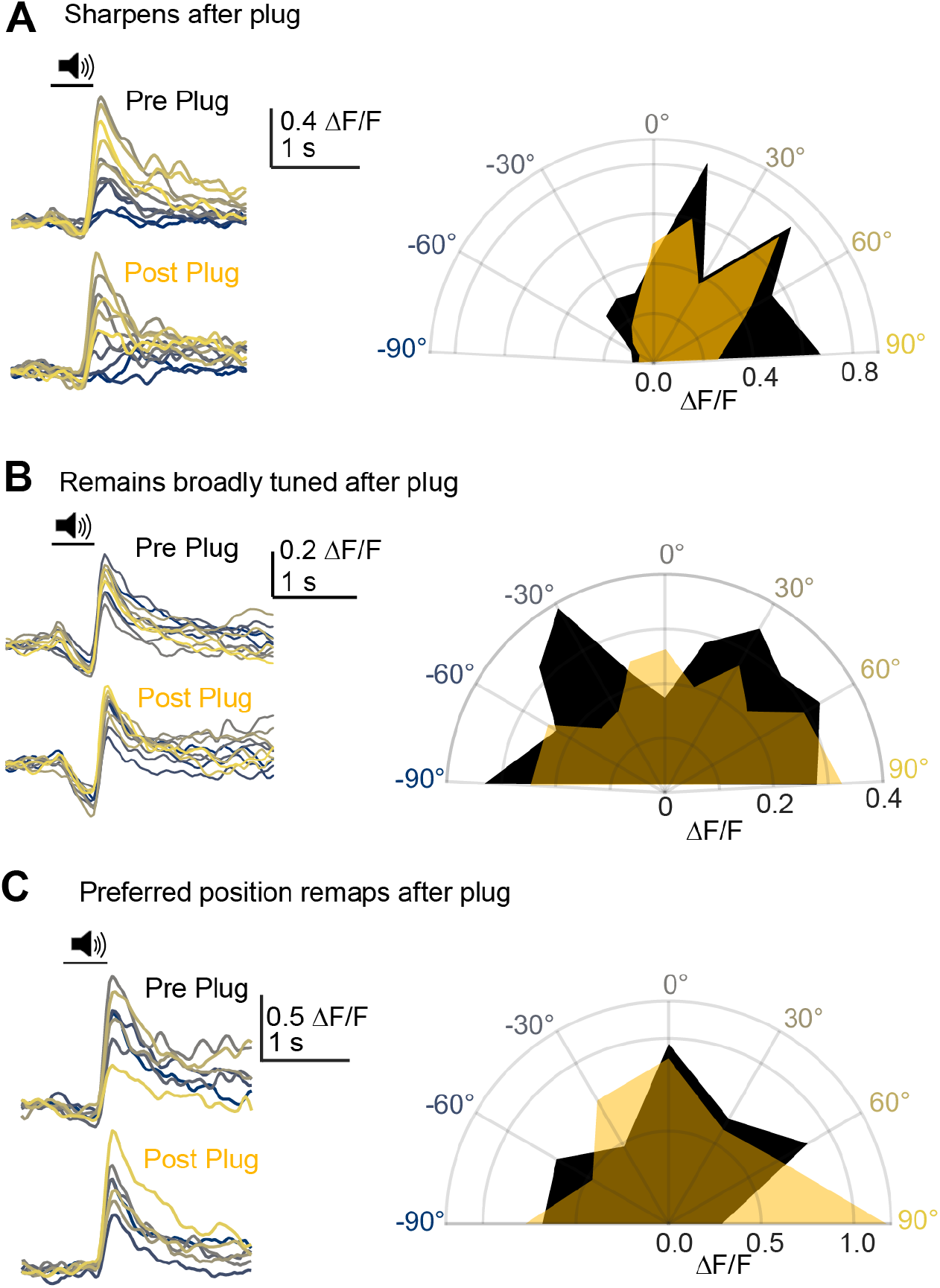
Additional examples of Group 2 neurons before and after plugging. A-C) Example average fluorescence traces and half-polar plots for group 2 neurons that sharpen (A), remain broadly tuned (B), or remap (C) after ear-plugging.

**Figure S6:**
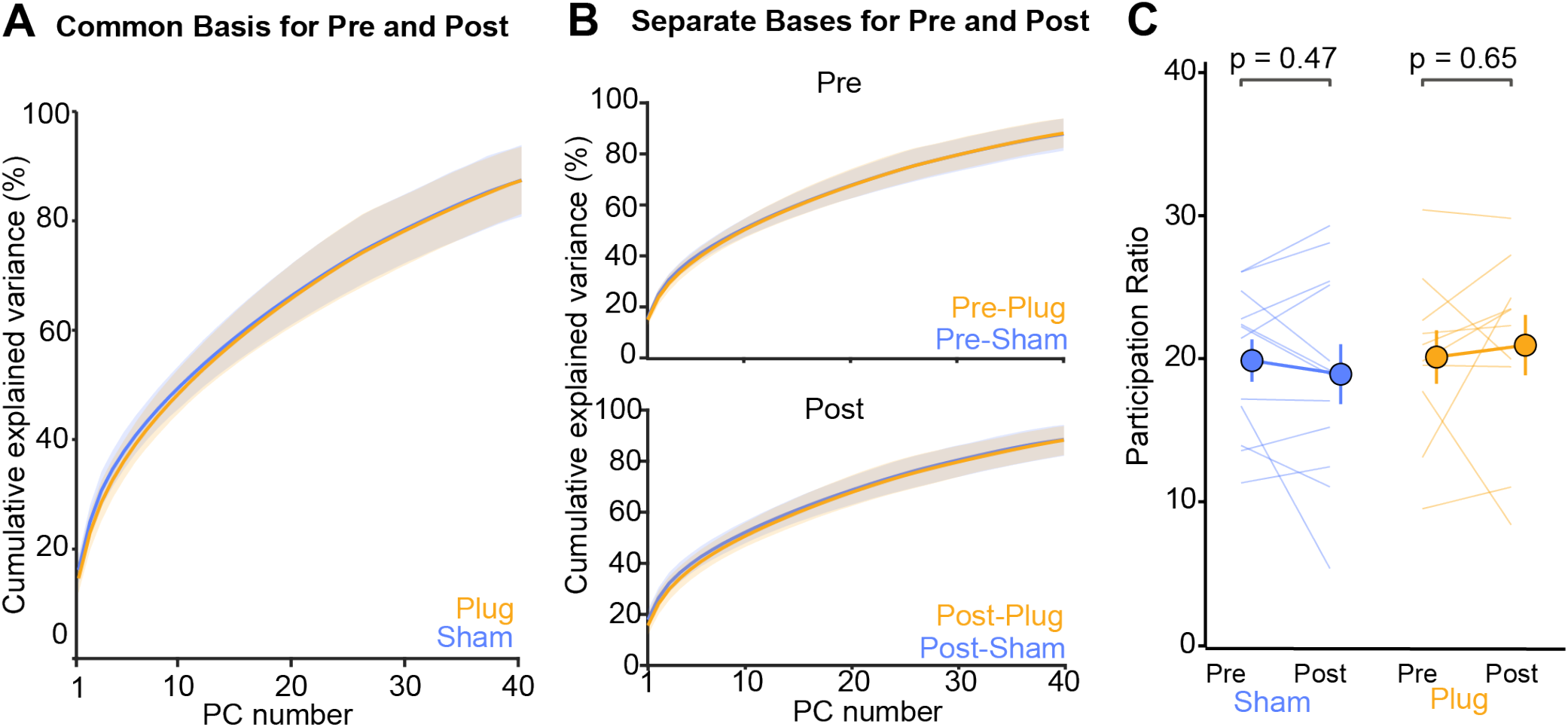
Ear-plugging does not alter effective population dimensionality. A) Cumulative variance explained as a function of principal component number, for common basis discovered using single trial data concatenated across pre- and post-treatment conditions of the same neurons. Orange and blue lines are averages across n = 10 and n = 12 plug and sham experiments, respectively; shading is 95% CI of mean, calculated from 1000 bootstraps across experiments. B) Cumulative variance explained when the PCA basis is discovered separately for pre- and post-treatment conditions. Colors are the same as (A). C) Effective dimensionality, calculated as the participation ratio, is plotted for pre- and post-treatment conditions in sham and plug cohorts. Thin lines are individual subjects; bubbles are mean ± SEM. P-values are from pairwise t-tests comparing pre- and post-conditions in the same treatment condition.

**Figure S7:**
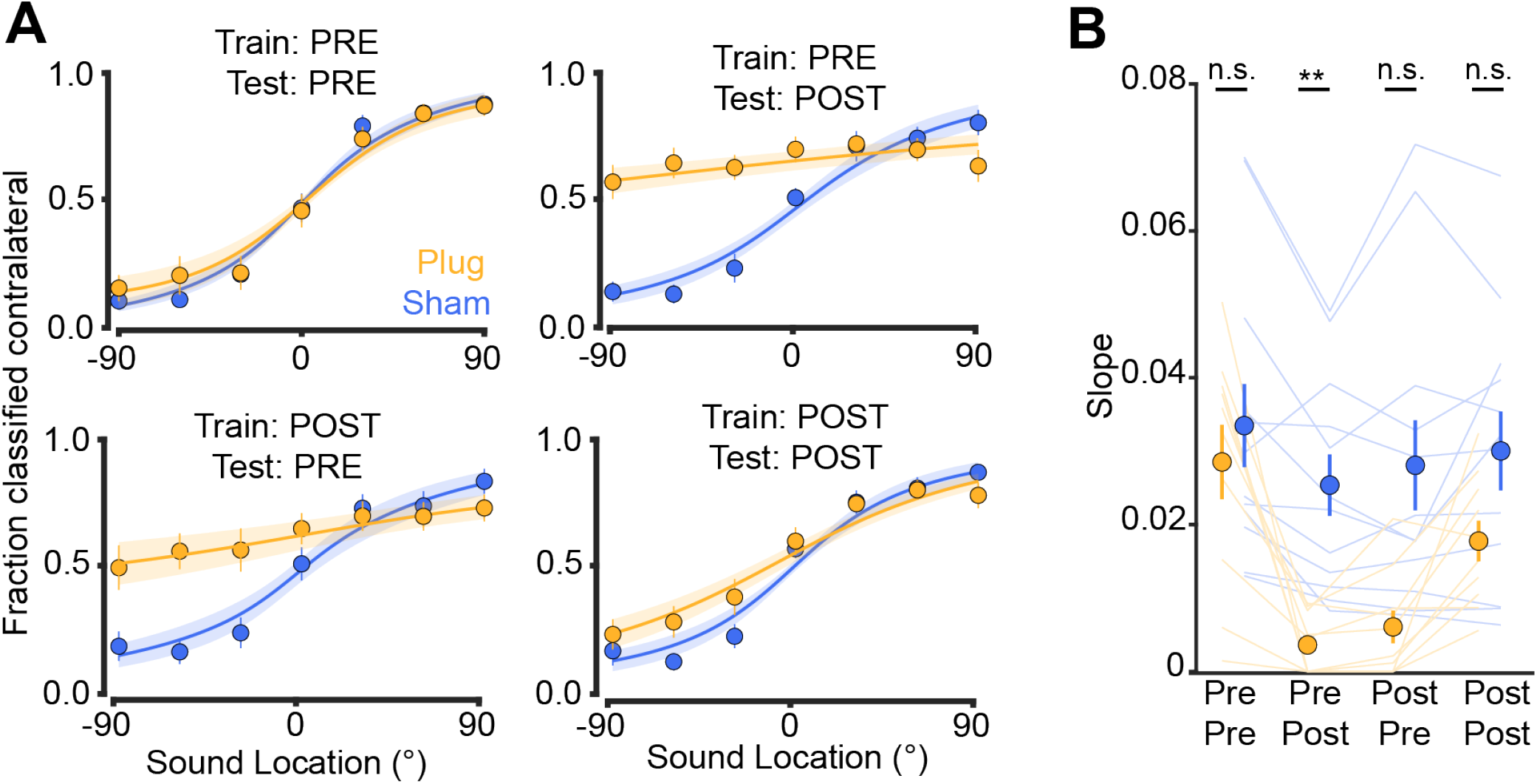
Psychometric evaluation of SVM classifiers. A) The fraction of trials classified as contralateral by SVMs trained and tested on different combinations of datasets. Results are shown for plug and sham treated cohorts in yellow and blue, respectively. Symbols are mean ± SEM, lines and shading are mean ± SEM of logistic fits to individual mice. B) The slope of the logistic fits in each condition are plotted for individual mice (lines) and for the population mean ± SEM (symbols). A two-way, mixed design ANOVA revealed main effects of treatment condition (F(1,20) = 6.37, p = 0.02), classifier training/testing (F(3,60) = 22.5, p < 0.001), and a treatment condition x classifier training/testing interaction (F(3,60) = 6.6, p<0.001). Asterisks denote significance of post-hoc tests comparing sham and plug within the same training/testing condition (p = 0.002). n.s., not significant.

## Notes

### Competing Interest Statement

The authors have declared no competing interest.

### Summary of Updates

Analyses and interpretations have been revised

